# Brain Infiltrated Monocyte-Macrophages in a rat model of Temporal Lobe Epilepsy: Revisiting the Pro-Inflammatory Paradigm

**DOI:** 10.1101/2025.01.02.631083

**Authors:** Wanda Grabon, Nadia Gasmi, Anatole Lang, Anne Ruiz, Béatrice Georges, Victor Blot, Michaël Ogier, Sylvain Rheims, Fabrice P Navarro, Laurent Bezin

## Abstract

Neuroinflammation is central to temporal lobe epilepsy, yet the specific role of myeloid cells remains unclear. In status epilepticus (SE) models, circulating monocytes have been reported to infiltrate the brain, though distinguishing them from microglia remains challenging. Using a rat model, we traced infiltrating monocytes post-SE, to investigate their persistence, phenotypic evolution during epileptogenesis and contribution to neuroinflammation. By tracking phagocyted fluorescent nanoparticles and using CD68 immunohistochemistry, we confirmed that monocytes entered the brain in significant numbers 24 hours post-SE, after the inflammatory peak occurred (7h post-SE). Tracked up to 7 weeks, these cells adopted a microglia-like phenotype, contributed to the microglial scar and sustained low-grade inflammation during the chronic phase of epilepsy solely through their presence, as their expression of pro-inflammatory markers resembled that of non-activated microglia. Importantly, monocytes initially and transiently supported an anti-inflammatory response providing a unique opportunity to modulate neuroinflammation and potentially disrupt epilepsy progression, opening new avenues for therapeutic interventions.

## 1. INTRODUCTION

Neuroinflammation plays a critical role in the pathophysiology of temporal lobe epilepsy (TLE), influencing epileptogenic network formation post-injury, cognitive impairments, and the maintenance of spontaneous seizures (*1*–*4*). Both experimental models and TLE patients show evidence of blood-brain barrier (BBB) leakage (*5*–*7*) and monocyte infiltration, particularly within the hippocampus (*8*– *11*). Initial studies of these infiltrating monocytes suggested that they play a major role in the early extensive secretion of pro-inflammatory cytokines after pro-epileptogenic brain aggression, such as *status epilepticus* (SE), and that they may play a detrimental role in the development of TLE (*10*–*12*). However, the fate of these infiltrating monocytes and their role in sustaining inflammation remain debated (*8, 10*). Through non-invasive tracking studies—crucial both for minimizing the animal suffering caused by SE induction procedures and for maintaining a physiological approach—and by controlling post-mortem tissue inflammatory responses (*13*), we provide new insights into the role of myeloid cells in post-SE inflammation and in established epilepsy.

Detecting infiltrating monocytes in the brain presents challenges due to their shared embryonic origin with microglia, resulting in near-identical phenotypes (*14, 15*), and their potential evolution into brain monocyte-macrophages (mo-mΦs) (*16, 17*). To accurately track monocyte entry and fate, monocyte markers must be both specific to monocytes and sustained over time. Markers such as CCR2, CX3CR1, and Ly6C have allowed differentiation between microglia and infiltrating monocytes at early post-SE stages in mice (*8, 10*–*12, 18*). However, the role of infiltrating monocytes has received much less attention in rat models of SE, predominantly due to the lack of reliable markers. Moreover, while monocyte infiltration has been documented for weeks or even months in models of multiple sclerosis (*19*), spinal cord injury (*19*), and Alzheimer’s disease (*20*), their long-term fate post-SE remains unexplored.

This study aimed to elucidate the fate of peripheral monocytes infiltrating the brain following SE induced by pilocarpine in young adult rats. By using circulating monocyte depletion strategies, fluorescent latex bead labeling, and fluorescence-activated cell sorting (FACS), we tracked the fate and persistence of monocytes during the chronic epilepsy phase up to 7 weeks post-SE. Additionally, we used immunodetection of CD68, a marker specific to mo-mΦs in rats (*21, 22*) and humans (*23*) at the histological level, to monitor their presence and phenotypic changes. A secondary objective was to distinguish the respective contribution of microglia, infiltrating monocytes and mo-mΦs in the inflammatory response from the onset of epileptogenesis to the chronic phase of epilepsy. We assessed the inflammatory profiles of microglia, monocytes, mo-mΦs, and other brain cells, quantifying inflammatory and neuroprotective marker transcripts using RT-qPCR while minimizing *ex vivo* transcription and translation during the dissociation and sorting process (*13, 24*).

## 2. RESULTS

### 2.1 Myeloid cell reactivity in the hippocampus following pilocarpine-induced SE

In addition to the rapid microglial response observed following SE, pioneering studies have shown that monocytes infiltrate the brain parenchyma. However, the long-term fate of these infiltrating cells has not yet been established. CD11b immunodetection enables a detailed assessment of myeloid cell morphology, reflecting their state of activation and providing an indication of their phenotype, as illustrated in the whole hippocampus from 7h to 7 weeks post-SE (**Fig. S1**). In slices from healthy control rats, resting CD11b-positive microglia displayed a ramified morphology. Rapidly (7h) after SE, almost all microglial cells were in a reactive state, with enlarged cell body and retracted ramifications. At one day post-SE, monocyte-like round CD11b cells, namely in the hilus of dentate gyrus (DG), could be distinguished from reactive bushy cells (arrowheads, **Fig. S1F**). At this stage, no CD11b-positive cells recovered a resting morphology in the hippocampus. Six days post-SE, a CD11b+ hypersignal, which we will refer to as microglial-like scar, was clearly visible in the hilus of the DG and CA1 (**Fig. S1G-H**). however, this signal persisted predominantly in CA1, 7 weeks post-SE during the chronic phase of epilepsy (**Fig. S1I-J**), while CD11b-positive microglial cells returned to their basal morphological state in other areas of the hippocampus. This scar could play an adaptive role by containing inflammatory damage and preserving the integrity of the surrounding tissue. The prolonged persistence of microglial activation in the CA1 region, compared to the DG, could suggest region-specific differences in the ability of microglia to regulate tissue damage in relation to the resolution of inflammation. It is also possible that this different level of CD11b activation may be a signature of a distinction between resident microglial cells and myeloid cells of peripheral origin, which may have established themselves more durably in certain regions rather than others.

### 2.2 Peripheral origin of round-shaped CD11b-positive cells

The presence of CD11b-positive round cells 24h post-SE coincides with the early transient induction of chemokines monocyte chemoattractant protein 1 (MCP-1) and macrophage inflammatory protein α (MIP1α) in the hippocampus (**Fig. S2**). This suggests that many of these round cells are likely infiltrating monocytes, although we cannot entirely rule out the possibility that some might be amoeboid microglia. To demonstrate the peripheral origin of round CD11b-positive cells and to track them as long as possible in the brain tissue after SE, we tagged peripheral monocytes prior to the induction of SE. We first depleted circulating monocytes/macrophages (*25*) 3 days before the induction of SE by administrating clodronate liposomes. We then administered intravenously FYG nanoparticles to label newly generated circulating monocytes (**Fig. 1A**). We used CD11b marker to detect myeloid cells, as at the onset of this study, no specific marker was available in rats to reliably distinguish microglial cells from infiltrating monocytes *in situ*. Detection of CD11b and FYG was performed at 1-, 3- and 6-days post-SE in the hilus and provided evidence that infiltrating monocytes remained in the hippocampus for at least 6 days post-SE. We observed that at 1-day post-SE, CD11b-positive round-shaped cells located in the blood vessels, adjacent to the vascular wall, were also positive for FYG (**Fig. 1B**). At 3 days post-SE, fluorescent nanoparticles were found in CD11b-positive cells which morphology was similar to that of activated microglial cells (**Fig. 1C**). At 6 days post-SE, FYG nanoparticles were still observed in CD11b-positive cells (**Fig. 1D**) but at a lower density, which may be explained by the fact that the CD11b-positive cells that had phagocytosed the FYGs may have processed or degraded them. It is noteworthy that fluorescent nanoparticles have not been detected in Iba1-positive cells (data not shown). It is therefore likely that at 6 days, the CD11b-positive cells labeled with FYG were cells resulting from the evolution of monocytes into brain mo-mΦs. The condition of rats treated with clodronate prior to the induction of SE deteriorated beyond that of rats subjected to SE alone, reaching the predefined endpoints 6 days post-SE, which prevented the continuation of the study beyond this time.

**Figure 1.**
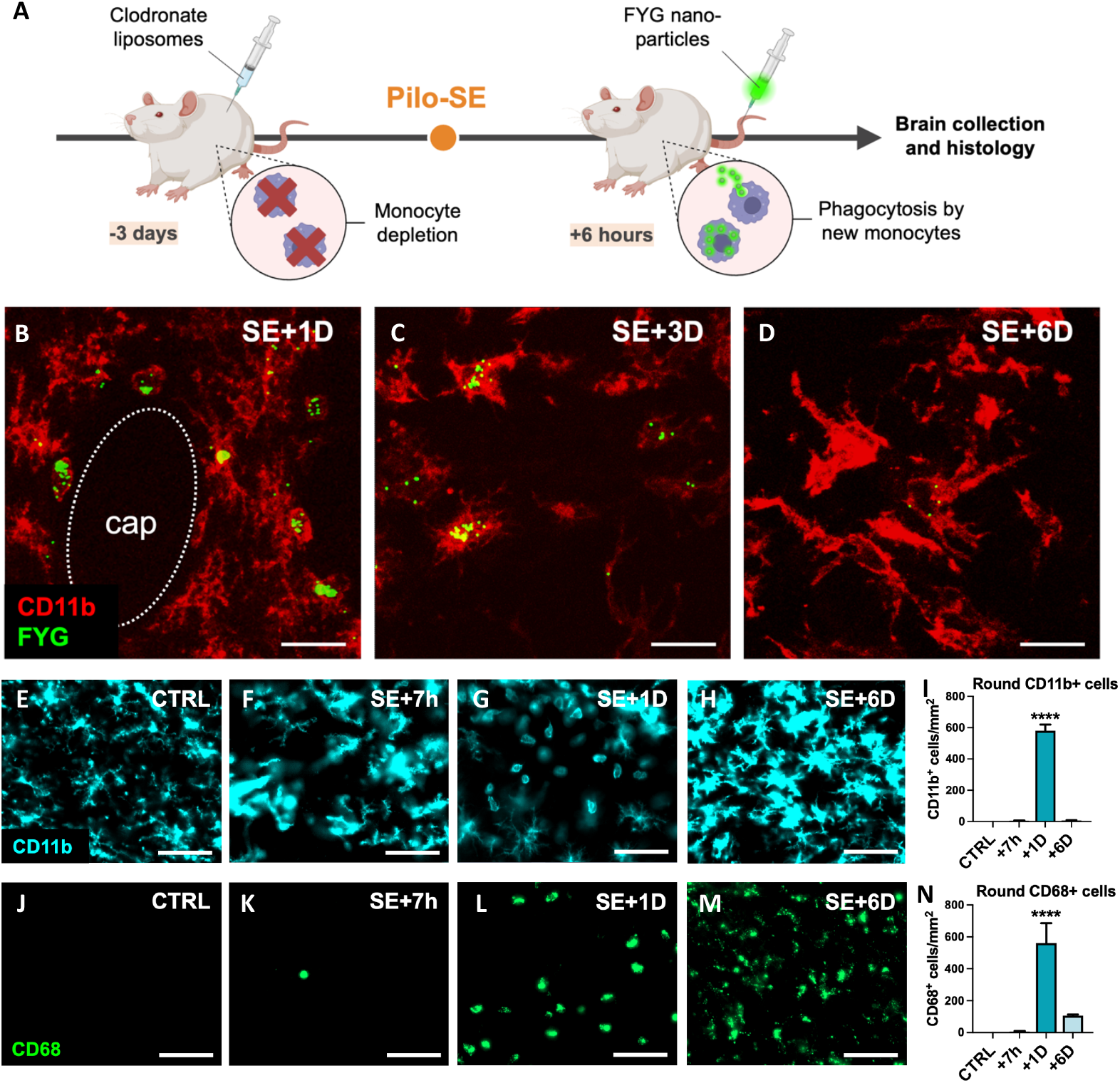
Peripheral monocytes infiltrate the hippocampus following SE between +7h and +6 days and differentiate into brain monocyte-macrophages. **A-D**. Fluoresbrite YG carboxylate microspheres (FYG, 0.5 μm) were injected to the tail vein 6h post-SE. No labeled monocyte could be detected in the brain parenchyma unless circulating monocytes were depleted with clodronate liposomes (1 mL/100g; i.p.) administered prior to SE. Rats were sacrificed 1D, 3D and 6D post-SE. Detection of CD11b (Red, CBL1512Z, Millipore) and FYG (green) in infiltrating monocytes at 1 day (**B**, cap=capillary), brain monocyte-macrophages with extending processes at 3 days post-SE (**C**) and in cells resembling activated microglia 6 days post-SE (**D**) in the hilus. Scale: 20 µm. **E-N**. CD11b (**E-I**, cyan, CBL1512Z, Millipore) and CD68 (**J-N**, green, MCA341GA, Bio-rad) were immunodetected in the dentate gyrus following SE (CTRL, n=6; SE+7h, n=4; SE+1D, n=5; SE+6D, n=4). Scale : 50µm. Round CD11b-positive cells (**J**) and CD68-positive cells (**N**) were quantified in the dentate gyrus. Data are analyzed with Tukey’s test following one-way ANOVA. Data are presented as mean + SEM. *: vs. CTRL. ***, p<0.001; ****, p<0.0001.

The CD68 marker has recently emerged histologically as a specific marker of monocytes in rats, particularly in epileptogenic situations (*21*), which we confirmed here using Iba1 and CD68 co-staining (**Fig. S3**). Unfortunately, the time elapsed between conducting the experiment on clodronate-FYG rats and the identification of CD68 as a monocyte-marker was too long, leading to a complete degradation of FYG fluorescence, and rendering CD68-FYG co-detection impossible. Nevertheless, when numerous round CD11b-postive cells were detected 24h post-SE in the hippocampus, numerous round CD68-positive cells were also detected in the adjacent section, in similar number (**Fig. 1E-N**). No or rare CD68-positive cells were detected in control brain sections, or 7h post-SE, respectively (**Fig. 1J,K**). The number of CD11b-positive round cells quantified in the dentate gyrus between 7h and 6 days post-SE is strongly associated with the number of CD68-positive round cells (simple linear reagression, p<0.0001, R^2^=0.8043,y = 0.9255x + 40.11).

### 2.3 Some monocyte-macrophages persist in the hippocampus in the long-term, once epilepsy has developed

We tracked the presence of infiltrated monocytes up to 7 weeks post-SE using CD68, and monitored their evolution into microglial-like cells based on morphology and co-labeling with the microglial marker Iba1 (**Fig. 2**). CD68-positive round cells were predominantly found in the DG and CA1, with a likely shift in peak entry between the two regions (**Fig. 2K,L**). Only a few CD68-positive round cells were detected 7 weeks post-SE, once epilepsy was established. Nine days after SE, CD68 colocalized with Iba1 in some non-round cells (**Fig. 2D,I**), supporting the hypothesis that CD68-positive monocytes infiltrating the brain parenchyma post-SE began to evolve into microglia-like cells. These morphologically microglia-like cells, both CD68-positive and Iba1-positive, were still detectable 7 weeks after SE (**Fig. 2E,J**), during the chronic epilepsy phase, notably in CA1. In this manuscript, we will refer to them as ‘monocytes’ upon entry, up to the 24-hour time point, and then as monocyte-macrophages (mo-mΦs) at later time points, once their phenotype has evolved.

**Figure 2.**
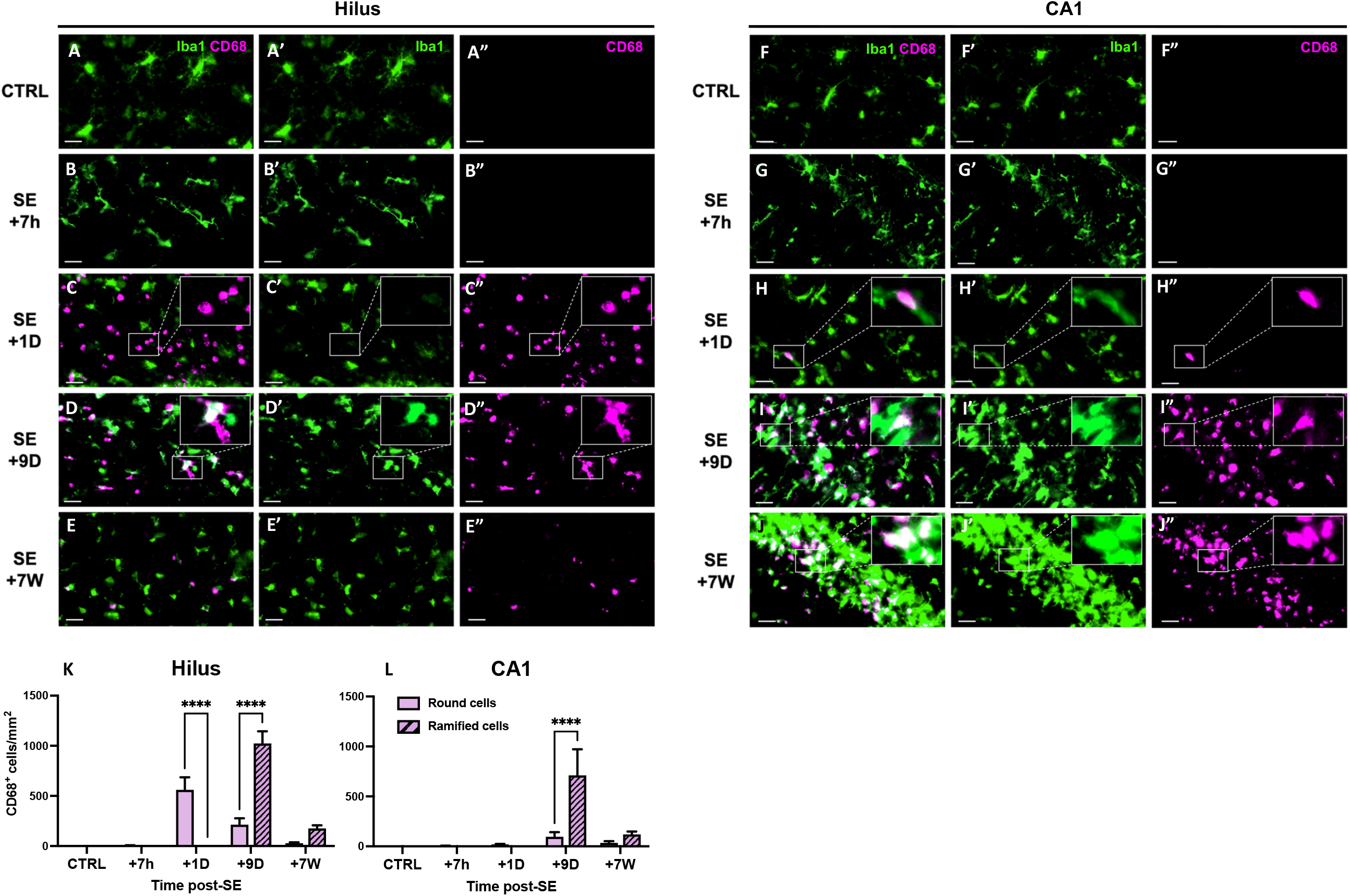
Infiltrating monocytes differentiate into brain monocyte-macrophages bearing morphological feature of microglial cells. **A-J**. CD68 (MCA341GA, Bio-rad) and Iba1 (ab5076, abcam) were immunodetected in the dentate gyrus (left) and CA1 region (right), in control rats and 7h, 1D, 9D and 7W post-SE. Images were acquired with a slide scanner, objective ×20. Scale bars: 20µm. **K-L**. Quantification of round and ramified CD68-positive cells in the hilus and in CA1 region, CTRL, n=5; SE+7h, n=4; SE+1D, n=5; SE+9D, n=4; SE+7W, n=5. Data are analyzed with Tukey’s test following two-way RM ANOVA. Data are presented as mean + SEM. **** p<0.0001. Only significant differences between round and ramified cells are shown. All statistical tests are detailed in table S2.

### 2.4 Infiltration of monocytes into other vulnerable regions subjected to massive neuroinflammatory processes after SE

Brain-infiltrating monocytes were observed not only in the hippocampus, but also in the ventral limbic region (VLR), which includes (*26*) the amygdala, the piriform cortex, and the agranular insular cortex, as well as in the dorsal thalamus (ThD). They infiltrate these regions within 24h post-SE, following the induction of chemokine MCP1, and subsequently evolve into mo-mΦs, as described earlier for the hippocampus (**Fig. S4**). It is noteworthy that monocytes tend to accumulate in areas where the so-called ‘microglial scar’ is present, typically visualized using CD11b staining (**Fig. S4A-D**). This strongly suggests that the infiltrating monocytes remain in the brain parenchyma and transform into activated microglia-like monocyte-macrophages, which very likely constitute the cells that form the microglial scar (**Fig. S4**).

To determine the respective contribution of microglia, and monocytes/mo-mΦs and other non-myeloid cells to the inflammatory response, we first characterized the tissue inflammatory status at the transcriptional level using RT-qPCR on homogenized brain tissue samples. We then investigated these inflammatory markers in FACS-sorted cell populations. To increase cell yield and optimize the likelihood to obtain a sufficient number of cells, particularly at the +7 weeks timepoint for subsequent RT-qPCR studies, sorting was performed on pooled samples from the three regions we identified with the highest monocyte infiltration: the hippocampus, the VLR, and the ThD. To ensure comparability between tissue and sorted cell results, we present tissue-level quantifications as the average of the three regions (**Fig. 3**), with detailed structure-specific data shown in **Figure S5**.

**Figure 3.**
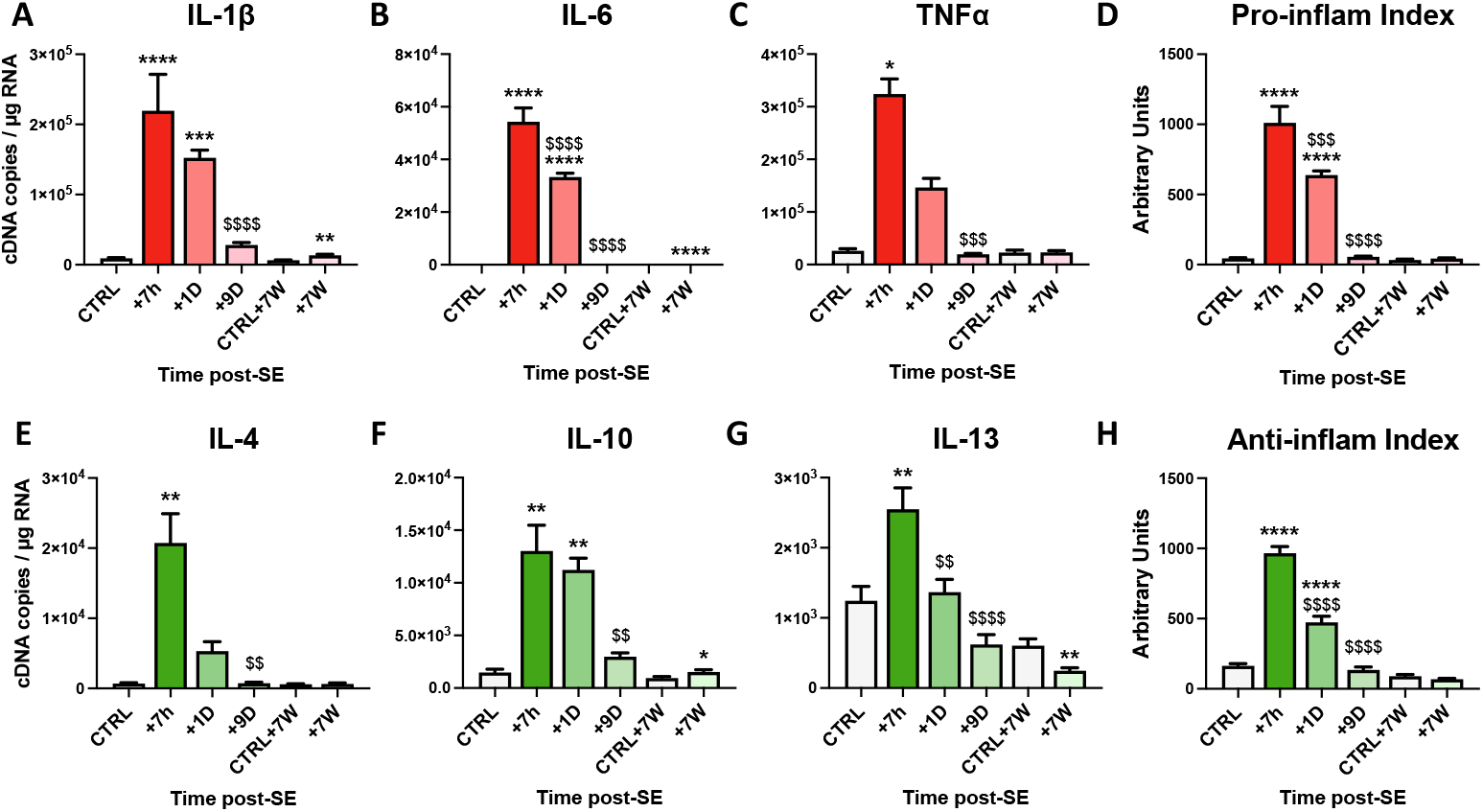
Transcriptional inflammatory response in the hippocampus, the VLR and the ThD during epileptogenesis and chronic epilepsy following SE induced by pilocarpine at P42 in rats. Prototypical cytokine mRNA levels were quantified using calibrated RT-qPCR in hippocampi, ventral limbic region (VLR) and dorsal thalamus (ThD) following SE (CTRL, n=6; SE+7h, n=6; SE+1D, n=6, SE+9D, n=7, CTRL+7W, n=6, SE+7W, n=8). Transcript levels of each gene were measured in each structure, then averaged for each individual and expressed as cDNA copies per µg of total RNA. Levels by structure are detailed in Figure S5. Pro-inflammatory index (PI-I) and anti-inflammatory index (AI-I) were calculated as described in the Method section from IL-1β, IL-6 and TNFα and IL-4, IL-10 and IL-13 respectively, and expressed in arbitrary units. Data measured 7h, 24h and 9 days post SE were compared to P42 healthy controls, and those measured 7 weeks post SE were compared to another age-matched control group. Details of statistic tests are presented Table S2. All data are presented as mean + SEM. *: vs. respective CTRL; $: vs.SE+7h. */$, p<0.05; **/$$, p<0.01; ***/$$$, p<0.001; ****/$$$$, p<0.0001.

Results show a sharp but transient induction of transcript levels of pro-inflammatory cytokines IL-1β, IL-6 and TNFα that peaked 7h after SE onset (**Fig. 3A-C**, details of all the statistical tests are available in **Table S2**). Transcript levels of these pro-inflammatory cytokines were still higher than that quantified in healthy controls 24h after SE. IL-1β, IL-6 and TNFα transcripts were back to control levels 9 days following SE. However, IL-1β and IL-6 transcript levels were still significantly elevated (222.8 ± 35.2% and 117.1 ± 2.1% of respective controls, respectively) at 7 weeks post-SE during the chronic phase of epilepsy. Pro*-*inflammatory index (PI-I; see material and methods section) was calculated from these prototypic cytokine transcript levels to reflect overall pro-inflammatory response after SE (**Fig. 3D**). PI-I peaked at 7h post-SE and was still significantly higher from that of controls 24h post-SE. The inflammatory response was also accompanied by marked and transient expression of anti-inflammatory cytokines, as evidenced by elevated levels of IL-4, IL-10 and IL-13 transcripts 7h and 24h post-SE (**Fig. 3E-G**). As for the PI-I, an anti-inflammatory index (AI-I) was calculated from these prototypic cytokine transcript levels to summarize the general anti-inflammatory response after SE (**Fig. 3H**). AI-I peaked at 7h and 24h after SE and was back to control levels 9 days post-SE.

### 2.5. Microglial cells are major contributors to pro-inflammatory cytokine expression after SE

FACS was used to sort microglia (CD11b^+^CD45^lo^CD11a^lo^) and monocytes/mo-mΦs (CD11b^+^CD45^hi^CD11a^hi^)(*27, 28*), following CD11b-enrichment by MACS. CD11b-negative cells (remaining brain cells, i.e. neurons, astrocytes, oligodendrocytes, endothelial cells, etc.) were also collected (**Fig. 4A**). Hippocampus, VLR and ThD were microdissected and pooled for FACS studies, as brain regions where monocyte infiltration was strong (**Fig. S4**). The study was performed at different times post-SE, i.e. 24h, 9 days and 7 weeks post-SE. The entire tissue dissociation and cell sorting protocols of this study were performed in the presence of transcription and translation inhibitors in all buffers used to restrain *ex vivo* cell activation and obtain the most reliable inflammatory profile possible (*24*). In control rats, almost no CD11b^+^CD45^hi^CD11a^hi^ (monocytes/mo-mΦs) cells were detected (**Fig.S6**). The highest proportion of monocytes/mo-mΦs was quantified 24h post-SE.

**Figure 4.**
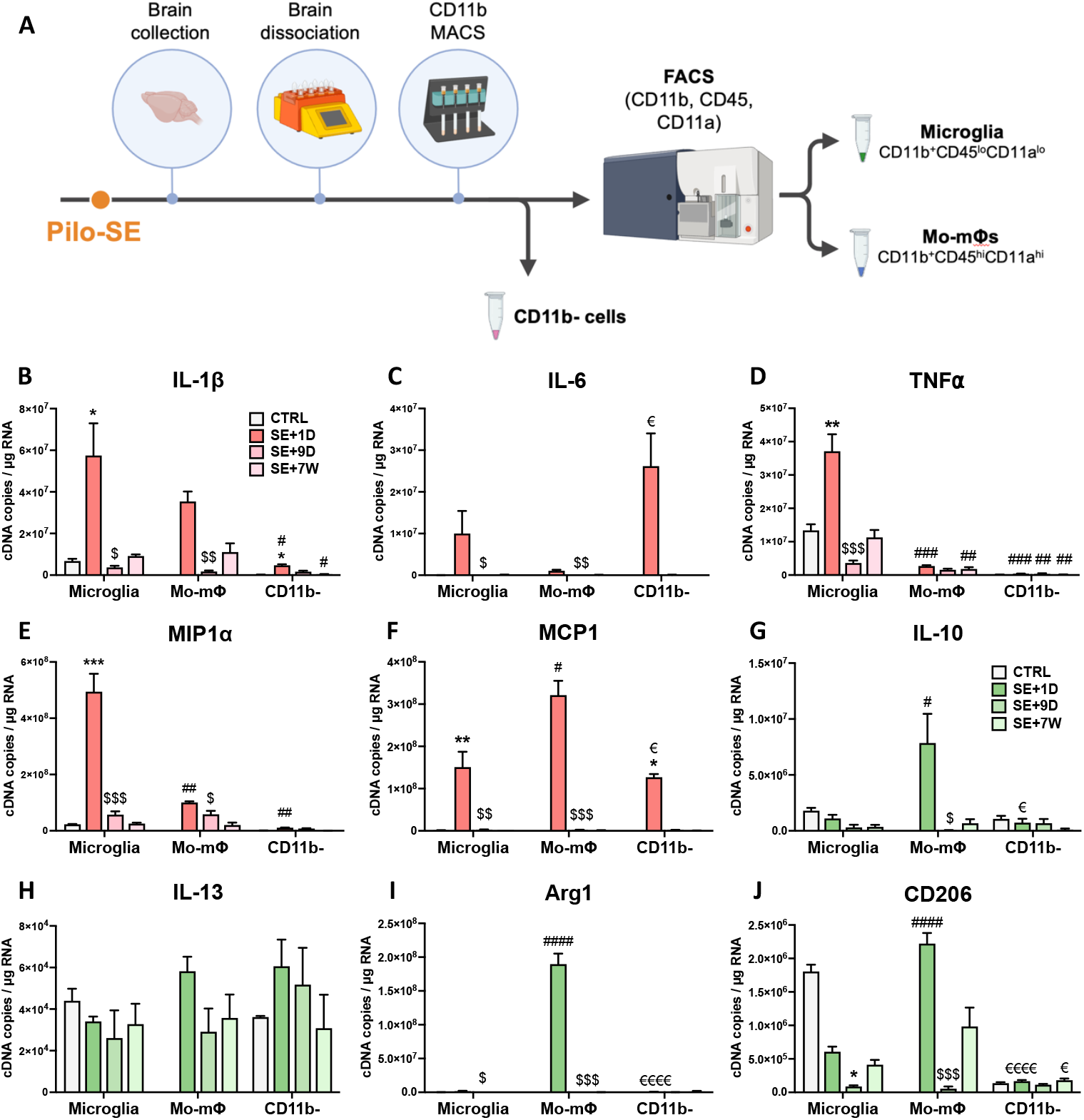
Transcriptional differences between sorted microglia, monocytes/mo-mΦ and CD11b-negative cells after SE. **A**. CD11b-cells, microglia and monocytes/mo-mΦ were sorted by MACS and FACS. **B-J**. Pro-inflammatory cytokines (IL-1β, IL-6, and TNFα), chemokines (MIP1α, MCP1) and immunomodulatory markers (IL-10, IL-13, Arg1, CD206) transcript levels were quantified by RT-qPCR in sorted cell populations (CTRL, n=2; SE+1D, n=3; SE+9D, n=3; CTRL 7W, n=3; SE+7W, n=3). Data measured 7h, 24h and 9 days post SE were compared to P42 healthy controls (“CTRL”), and those measured 7 weeks post-SE were compared to another age-matched control group (“CTRL+7W”). Details of statistic tests are presented in Table S2. Results are presented as mean + SEM. “Time post-SE” factor: *, vs. respective CTRL; $, vs. respective SE+1D; “cell type” factor: #, vs. respective microglia; €, vs. respective mo-mΦ. 1 symbol, p<0.05; 2 symbols, p<0.01; 3 symbols, p<0.001; 4 symbols, p<0.0001.

Furthermore, in line with what we observed at the histological level, we report the long-term presence of brain mo-mΦs in the epileptic tissue, their proportion being still significantly above that measured in controls 7 weeks after SE (**Fig. S6F**).

We quantified the transcript levels of the myeloid cell markers CD11a and CD11b, whose corresponding proteins were used for cell sorting to distinguish microglia (CD11b^+^CD11a^lo^) from monocytes/mo-mΦs (CD11b^+^CD11a^hi^), using RT-qPCR on the sorted cell populations. Addiionally, we measured the transcript levels of the microglial marker Iba1 and of the monocyte marker CD68. Although we anticipated CD11b mRNA to be expressed in both microglia and monocytes, we expected Iba1 mRNA to be restricted to microglia and CD11a and CD68 mRNAs to be exclusive to monocytes/ mo-mΦs. However, this was not the case, as all four mRNAs were detected at nearly similar levels in both cell populations, with only occasional statistically significant differences (**Fig. S7**).

Transcript levels of the 3 prototypical pro-inflammatory cytokines (IL-1β, IL-6 and TNFα), 2 chemokines (MIP1α and MCP1) and 4 immunomodulatory-associated genes (IL-10, IL-13, Arg1 and CD206) were quantified by RT-qPCR in the sorted cell populations from 24h, i.e., once monocytes have infiltrated the brain, to 7 weeks post SE (**Fig. 4B-J**).

IL-1β transcript levels were significantly higher in microglia sorted from brains collected 24h after SE than in control microglia (**Fig. 4B**). At this timepoint, IL-1β transcript levels were also detected at comparable levels in infiltrating monocytes and were slightly induced in CD11b-negative cells. IL-1β transcript levels were back to basal levels 9 days post-SE. The highest levels of IL-6 transcript were detected in CD11b-negative cells 24h after SE (**Fig. 4C**). These were significantly higher than those measured in infiltrating monocytes, but not than those measured in microglia. TNFα transcript levels were increased in microglia sorted from brains collected 24h after SE compared to control microglia, and were significantly higher than those measured in monocytes, with copy numbers almost 14-fold higher in microglia (**Fig. 4D**). At 7 weeks post-SE, TNFα transcript levels were still 6-fold higher in microglia than in mo-mΦ. At each timepoint studied, levels were higher in microglia than in CD11b-negative cells. The transcript level of the inflammatory chemokine MIP1α was significantly increased at 24 hours in microglia, reaching a level 5 times higher than in mo-mΦ (**Fig. 4E**). The mRNA level of MCP1, a chemokine primarily involved in monocyte recruitment, was strongly and transiently induced 24h post-SE across all three sorted cell populations, with the highest level measured in monocytes (**Fig. 4F**).

Overall, microglia appear to be the most pro-inflammatory cells during the intense inflammatory response observed during epileptogenesis, returning to a baseline level during the chronic phase of epilepsy. Monocytes made a smaller contribution to the expression of pro-inflammatory cytokines during epileptogenesis, primarily by expressing IL-1β 24h after SE induction. Mo-mΦs that remained in the tissue at 7 weeks exhibited low levels of IL-1β transcript expression. Consequently, the residual low-grade tissue inflammation observed at 7 weeks post-SE might be sustained by mo-mΦ that have become integrated into the brain parenchyma over the long term.

### 2.6. Brain-infiltrating monocytes exhibit an anti-inflammatory phenotype upon entry

Regarding anti-inflammatory cytokines, their significant induction at the tissue level 7 hours post-SE (**Fig. 3E-G**) cannot be attributed to infiltrating monocytes, as these cells only appear in brain tissue after this timepoint. By 1 day post-SE and beyond, their presence in brain tissue prompted an essential investigation into how the anti-inflammatory response is partitioned among microglial cells, mo-mΦs, and other brain cells. IL-10 transcript levels remained stable in microglia and in CD11b-negative cells following SE, but were high in infiltrating monocytes 24h following SE, suggesting that infiltrating monocytes are the main contributors of the increase in IL-10 transcripts observed at the tissue level 24h post-SE (**Fig. 4G**). IL-13 transcript levels remained also stable in microglia and in CD11b-negative cells following SE, and were detected in monocytes/mo-mΦs at similar levels (**Fig. 4H**). Transcript levels of Arg1, a characteristic marker of “M2” anti-inflammatory phenotype, was detected at dramatically higher levels in infiltrating monocytes 24h post-SE than in microglia and CD11b-negative cells (**Fig. 4I**). Transcript levels of immunomodulator-marker CD206 decreased following SE in microglia. At 24h post-SE, transcript levels of CD206 were ∼ 4-fold higher in infiltrating monocytes than in microglia (**Fig. 4J**). Taken together, these results suggest that monocytes play a significant role in the establishment of the anti-inflammatory response upon their entry 24h post-SE, transiently adopting a neuroprotective phenotype.

## 3. DISCUSSION

The role of infiltrating monocytes in the robust inflammatory response following epileptogenic insults, such as SE, in comparison to that of resident microglia, as well as their long-term fate within brain tissue, remains a subject of ongoing debate.

In P42 rats, pilocarpine-induced SE triggered a strong inflammatory reaction not only in the hippocampus but also more broadly in the ventral limbic and thalamic regions of the brain within hours. At the cellular level, the inflammatory response was characterized by the early transient activation of microglia. Peripheral monocytes infiltrated the hippocampus in significant numbers between 7h and 9 days after the onset of SE. Their number peaks at 24h, but some remained detectable for several weeks post-SE. Monocyte engraftment and their evolution into brain mo-mΦs was evidenced histologically over the long term, with a phenotype resembling that of microglia. We demonstrated that mo-mΦs play a significant role in the formation of the glial scar in hippocampal subregions, particularly the CA1 region and the DG. This scar is a well-known characteristic of hippocampal sclerosis, commonly observed in various experimental models and patients with TLE, and had long been believed to be primarily composed of resident microglia (*29, 30*). At the molecular level, while neuroinflammation largely resolves within a few days, a low-grade, persistent inflammation is observed during the chronic phase of epilepsy. A massive activation of both pro- and anti-inflammatory ges was detected at the mRNA level. IL-1β and IL-6 transcript levels were still significantly elevated up to 7 weeks post-SE during the chronic phase of epilepsy. Data collected with FACS indicated that microglia were the primary contributors to the early inflammatory peak, while monocytes exhibited a more prominent M2-type, anti-inflammatory and neuroprotective state than microglia upon their entry 24h after SE. However, once they evolved into mo-mΦs, they might be the main contributors to the persistent low-grade inflammation observed during chronic epilepsy.

Measuring the inflammatory profile of infiltrating monocytes is particularly challenging because distinguishing them from resident microglia involves a cell sorting step whose protocol, notably the tissue dissociation, induces significant transcriptional and translational changes *ex vivo* (*24, 31*). In a pilocarpine-SE mouse model, microglia from the hippocampus remained rather immune suppressed compared to myeloid infiltrates that displayed a strong inflammatory profile during the early phase of epileptogenesis (*11*). In a KA-induced SE model, IL-1β levels were similar in FACS-isolated microglia and mo-mΦs following SE, while mo-mΦs provided greater levels of TNFα than activated microglia (*10*). In our study, where we took care to minimize *ex vivo* cell activation, we demonstrated that at their peak infiltration, monocytes temporarily exhibit an overall anti-inflammatory and neuroprotective phenotype, contrasting with prior conclusions about their inflammatory state (*10, 11*). In the chronic epilepsy phase, when microglia have recovered a baseline state in terms of inflammatory cytokine expression and morphology, some mo-mΦs remain in the tissue and express inflammatory cytokines, albeit at a low level. Therefore, we hypothesize that these monocytes, because of their presence, and not because they express higher levels of pro-inflammatory markers than microglia, fuel the persistent low-grade inflammation observed once epilepsy is established.

To establish the peripheral origin of the round cells observed 24 hours after SE, confirming the entry of monocytes into the brain tissue, we employed a fluorescent tracing technique through intravenous administration of FYG particles (*25*). This approach requires the depletion of peripheral myeloid cells before SE onset, which means that at the time of SE, the monocyte pool is not completely replenished, potentially leading to a reduction in the number of infiltrating cells compared to the normal condition. This diminished number of infiltrating monocytes may account for the aggravated phenotype we observed in comparison to rats subjected to standard SE induction. This is in line with the observation that preventing monocyte infiltration resulted in dampened hippocampal neurodegeneration in a kainate-SE mouse model (*12*), and was detrimental in models of stroke (*32, 33*) and AD (*34*).

In our study, the extended observation of monocytes and subsequent mo-mΦs relied on tracking cellular markers, notably CD68 and CD11a. However, it is still possible that some monocyte subpopulations do not express these markers. Additionally, our data do not provide definitive insights into whether the decline in mo-mΦs numbers over time is attributable to alterations in the measured markers’ expression or actual cell elimination. Once a sufficiently specific marker of mo-mΦs is fully validated, the use of genetic models will allow to address reliably these questions by conducting fate mapping studies over the long term. Of note, while no CD68 and CD11a protein labeling was detected in the brain under control conditions, their transcripts were detectable in microglia making them unsuitable as reporter genes for the establishment of a transgenic model. However, Arg1 was found to be entirely monocyte-specific, but was transiently expressed, only upon their entry into brain tissue. Arg1 had also been shown to be specifically expressed by infiltrating myeloid cells in the CNS in models of spinal cord injury and multiple sclerosis (*19*). The Arg1 promoter could therefore be considered for inducible expression of a reporter gene up to 24h post-SE. Future single-cell screening analysis will allow for the identification of other monocyte-specific genes.

Investigation of differences in expression profiles between microglia and monocytes/mo-mΦs was not carried out at the single-cell level, but on whole populations, separated by FACS after CD11b-MACS enrichment. Thus, our results do not allow to ascertain whether all cells within a given population share the same inflammatory state or if they comprise multiple subpopulations exhibiting varying levels of activation, with the latter hypothesis being the more likely scenario, considering the heterogeneity of both microglia and monocytes (*35*–*37*).

Taken together, our findings have allowed us to propose a time-course depicting the contribution of microglia and infiltrating monocytes/mo-mΦs to inflammation following pilocarpine-induced SE in rats, from early epileptogenesis to chronic epilepsy (**Fig. 5**). Importantly, our results challenge the traditionally held view of mo-mΦs as major pro-inflammatory contributors in this context. By carefully limiting *ex vivo* cellular activation, we reveal that microglia, rather than mo-mΦs, are the primary drivers of the pro-inflammatory response. This repositioning of roles not only enhances our understanding of post-SE inflammation but also underscores the need to refine therapeutic strategies targeting these distinct cellular populations. Specifically, infiltrating monocytes emerge as promising targets, either by enhancing their anti-inflammatory phenotype during the early phase of epileptogenesis or by reactivating it once epilepsy is established.

**Figure 5.**
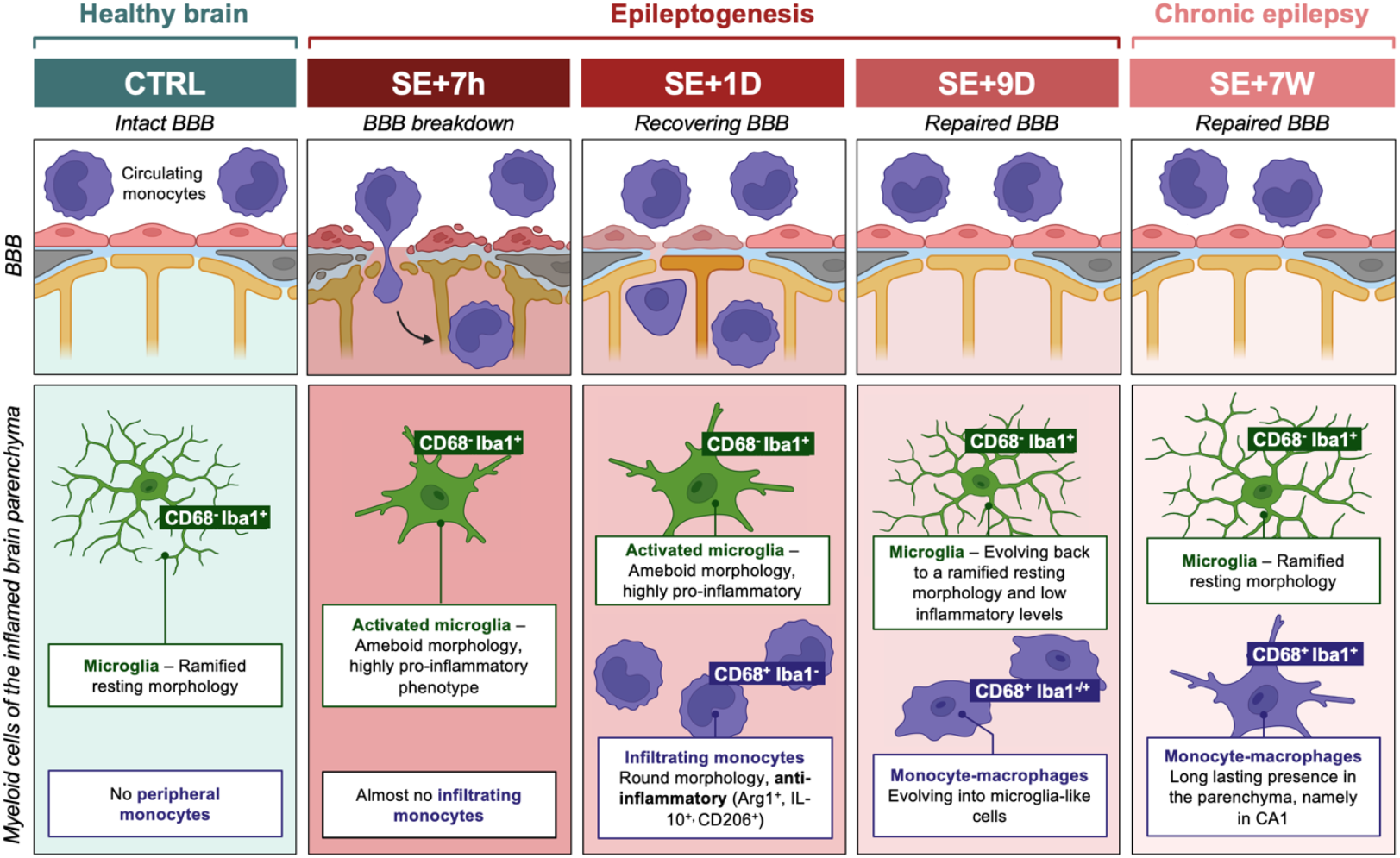
Proposed time-course of entry and integration of monocytes infiltrating the inflamed brain regions after pilocarpine-SE in rats. The blood-brain barrier (BBB) state following SE is depicted in the boxes at the top of the diagram (*38*). Myeloid cells in the hippocampus, the VLR and the ThD parenchyma, i.e. microglia and monocytes, are represented in the boxes at the bottom. Markers enabling the staining of both cell types at the histological level (but not at the transcript level) are highlighted in green for microglia and purple for monocytes/monocyte-macrophages (mo-mΦs). At 7 hours after SE, microglia are primarily responsible for the extensive expression of pro-inflammatory cytokines, and monocytes have not yet invaded the brain tissue. By 24 hours after SE, while inflammation remains at a significant level, a substantial number of round monocytes is observed, contributing to the expression of IL-1β but displaying a predominantly neuroprotective phenotype. Nine days after SE, a glial scar composed mainly of microglia and brain mo-mΦs forms, although the inflammatory response is largely resolved. At this stage, some mo-mΦs start to co express boss the monocyte-marker CD68 and the microglia-marker Iba1. During the chronic epilepsy phase, microglia have returned to a basal inflammatory state, but a low-grade inflammation persists, likely due to the presence of mo-mΦs. **Abbreviations**: BBB, blood brain barrier; CTRL, control, resting condition; D, day; SE, status epilepticus; W, week.

## 4. METHODS

Full details of the methods are given in the supplementary material.

### Experimental design

Four distinct groups of rats were similarly subjected to pilocarpine-induced SE at P42. Experimental design is illustrated in **Figure S8**.

#### Group 1 – Tracking of infiltrating monocytes following clodronate-induced depletion

Nine rats were used to track fluorescent-labeled monocyte infiltration following monocytes/macrophages depletion induced by clodronate administration. Brains were collected during epileptogenesis 1, 3 and 6 days following SE for subsequent immunohistology studies (SE+1D, n=3; SE+3D, n=3; SE+6D, n=3).

#### Group 2 – Detection of brain microglia, infiltrating monocytes and differentiated monocyte-macrophages during epileptogenesis and chronic epilepsy

Thirty-one rats were used to detect myeloid, microglial and monocyte markers using fluorescent immunohistochemistry during epileptogenesis 7h, 1 day, 6 days and 9 days following SE and during chronic epilepsy 7 weeks following SE in the hippocampus. CTRL, n=5; SE+7h, n=4; SE+1D, n=5, SE+6D, n=4 ; SE+9D, n=4; CTRL+7W, n=4; SE+7W, n=5.

#### Group 3 – Tissue inflammation at the molecular level during epileptogenesis and chronic epilepsy using RT-qPCR

Thirt-nine rats were used to evaluate inflammatory profiles at transcript level during epileptogenesis 7h, 1 day and 9 days following SE, and during chronic epilepsy, i.e. 7 weeks following SE in the hippocampus (CTRL, n=6; SE+7h, n=6; SE+1D, n=6, SE+9D, n=7; CTRL+7W, n=6; SE+7W, n=8).

#### Group 4 – Contribution of microglia and monocytes to neuroinflammation during epileptogenesis and chronic epilepsy using flow cytometry

Fourteen rats were used to evaluate inflammatory status of sorted microglia and monocytes/monocyte-macrophages using flow cytometry. Brains were collected during epileptogenesis 1 day and 9 days following SE and during chronic epilepsy, i.e. 7 weeks following SE for subsequent tissue dissociation and cell sorting (CTRL, n=3; SE+1D, n=3; SE+9D, n=3; CTRL+7W, n=2; SE+7W, n=3). Once sorted, the inflammatory profile of cell populations was evaluated by RT-qPCR.

### Animals

All animal procedures were in compliance with the guidelines of the European Union (directive 2010-63), taken in the French law (decree 2013/118) regulating animal experimentation, and have been approved by the ethical committee of the Claude Bernard Lyon 1 University (protocol # BH-2008-11). Male Sprague-Dawley rats (Harlan, France, and Envigo, France) were housed in a temperature-controlled room (23 ± 1°C) under diurnal lighting conditions (lights on from 6 a.m to 6 p.m) with water and food *ad libitum*.

### Pilocarpine-induced status epilepticus

SE was induced by pilocarpine, injected at P42. To prevent peripheral cholinergic side effects, scopolamine methylnitrate (1 mg/kg in saline, s.c.) was administered 30 min before pilocarpine hydrochloride (350 mg/kg, in saline, i.p.) (*39*). After 2h of continuous behavioral SE, diazepam (10 mg/kg, i.p.) was injected, followed 60 min later by a second injection of diazepam (5 mg/kg) to terminate behavioral seizures. Given the severity of the model, animals received special care as described in supplementary material.

### Clodronate-induced monocyte/macrophage depletion and fluorescent -nanoparticle labelling

Animals received clodronate liposomes (5mg/mL, i.p.) at the dosage of 1 mL per 100g body weight (*40, 41*). Clodronate was administered as a single dose 3 days before SE. Then, rats underwent SE and received 6 hours after its onset an injection of fluoresbite YG carboxylate microspheres (0.5 μm diameter) via the tail vein (9.1 × 10^10^particles per rat).

### Immunohistology

#### Brain slices

Forty μm-thick coronal sections were cut from frozen paraformaldehyde (PFA)-fixed brains using a cryomicrotome.

#### Fluorescent immunolabelling

Detection and morphology appreciation of myeloid cells was performed with mouse anti-Cluster of differentiation 11b (CD11b) antibody (1:2000, CBL1512Z, Millipore). Detection of microglia was performed with goat anti-ionized calcium binding adaptor molecule 1 (Iba1) antibody (1:500, ab5076, Abcam). Infiltrating monocytes and monocyte-macrophages (mo-MΦ) were labelled with mouse anti-CD68 antibody (1:1000, MCA341GA, Bio-rad). Fluorescent secondary antibodies are listed in supplementary material. Nuclei were stained with DAPI (300 nM, Molecular Probes).

#### Microscopy

Whole sections were scanned with a Carl Zeiss Axio Scan.Z1 Digital Slide Scanner (ZEISS) with a X20 lens on a 6µm stack, using the pilot Zen (ZEISS), or observed with a TCS SP5X confocal microscopy system (Leica). Images were processed on with Fiji software (ImageJ). Counting of round vs. branched cells was performed automatically using the ‘Analyze Particle’ function of Fiji software.

### Reverse transcription and real-time quantitative PCR

Tissues from perfused-brains were crushed using Tissue-Lyser II (Qiagen) according to the manufacturer’s instructions. Total RNAs from brain structures were extracted using Tri-Reagent LS. Total mRNAs were then reverse transcribed to complementary DNA (cDNA) using both oligo dT and random primers with PrimeScript RT Reagent Kit (Takara, #RR037A) according to manufacturer’s instructions in the presence of a synthetic external non-homologous poly(A) standard messenger RNA (SmRNA) to normalize the RT step, as previously described (*42*). Each cDNA of interest was amplified using the Rotor-Gene Q thermocycler (Qiagen), the SYBR Green PCR kit (Qiagen, #208052) and oligonucleotide primers (Eurogentec) specific to the targeted cDNA (**Table S1**). cDNA copy number detected was determined using a calibration curve, and results were expressed as cDNA copy number/µg tot RNA.

Pro-inflammatory index (PI-I) and anti-inflammatory index (AI-I) were calculated using a specific set of pro-inflammatory and anti-inflammatory genes: IL-1β, IL-6, TNFα and IL-4, IL-10, IL-13, respectively, using the formula given in the supplementary material (*13*).

### Flow cytometry

Samples were processed for tissue dissociation and cell sorting immediately following brain collection as quickly as possible, on ice. To prevent any artifactual *ex vivo* gene expression changes during brain dissociation and cell sorting procedures, all buffers and solutions used during the process (from animal perfusion to sorted cell flash freezing) were supplemented with an “inhibition cocktail” composed of Actinomycin D (3µM), Anisomycin (100 µM) and Triptolide (10 µM) i.e. transcription and translation inhibitors (*13, 24*).

#### Brain tissue dissociation

Once collected, tissues were cut in smaller pieces with a scalpel and processed for dissociation using Miltenyi’s Adult Brain Dissociation Kit (#130-107-677) according to manufacturer’s instruction, running program 37C_ABDK_01. The inhibition cocktail was added in each reagent. Following debris removal, cells were counted manually (with trypan blue) before magnetic sorting.

#### CD11b-positive cells magnetic enrichment

To increase Fluorescence-Activated Cell Sorting (FACS) yields and efficiency, cell suspensions were first enriched using the Magnetic-Activated Cell Sorting (MACS) technique, magnetically separating CD11b-positive cells (microglia and infiltrating monocytes) for subsequent FACS using CD11b/c MicroBeads according to manufacturer’s instructions (Miltenyi #130-105-634), from other cells before direct freezing. Details are provided in supplementary material.

#### FACS

Microglia (CD11b^+^CD45^lo^CD11a^lo^) and infiltrating monocytes (CD11b^+^CD45^hi^CD11a^hi^) were sorted with BD FACS Aria™ III Cell Sorter (BD Biosciences), as detailed in supplementary material. Gating strategy is presented in **Figure S9**.

### Statistical Analysis

Statistical analyses were performed using Prism 10.0 software (GraphPad, USA), as detailed in supplementary material. Results are presented as mean + SEM (standard error of the mean). Differences with a p-value<0.05 (p<0.05) were considered to be statistically significant. Details of statistic tests for each figure are presented in supplementary data (**Table S2**).

## Author contributions

Conceptualization and design: LB, FN, NG, WG

Methodology: LB, FN, NG, WG

Investigation: AL, FN, NG, WG, AR, BG, VB, MO

Visualization: LB, WG

Supervision: LB

Writing-original draft: LB, NG, WG

Writing-review & editing: FN, SR

## Data and materials availability

All data are available in the main text or in the supplementary materials.

