## Supplementary data for "Brain Infiltrated Monocyte-Macrophages in a rat model of Temporal Lobe Epilepsy: Revisiting the Pro-Inflammatory Paradigm"

**Figure S1 –** Evolution of glial cell activation in the hippocampus after pilocarpine induced-SE.

**Figure S2 –** Transcript levels of chemokines are induced early during epileptogenesis in the hippocampal tissue.

**Figure S3 –** CD68 allows for detection of infiltrating monocytes in rat hippocampus following SE.

**Figure S4 –** Monocyte infiltration is strong in both hippocampus, ventral limbic region and dorsal thalamus and contributes to the formation of the microglial scar.

**Figure S5 –** Transcriptional inflammatory response in the hippocampus, the VLR and the ThD during epileptogenesis and chronic epilepsy following SE induced by pilocarpine at P42 in rats.

**Figure S6 –** Sorting of microglia and monocytes/mo-mΦs up to 7 weeks after SE by FACS.

**Figure S7 –** Cellular markers transcript levels in sorted microglia, monocytes/mo-mΦ and CD11b-negative cells after SE.

**Figure S8 –** Experimental design.

**Table S1 –** Sequences of primer pairs used for qPCR.

**Table S2 –** Details of statistic tests.

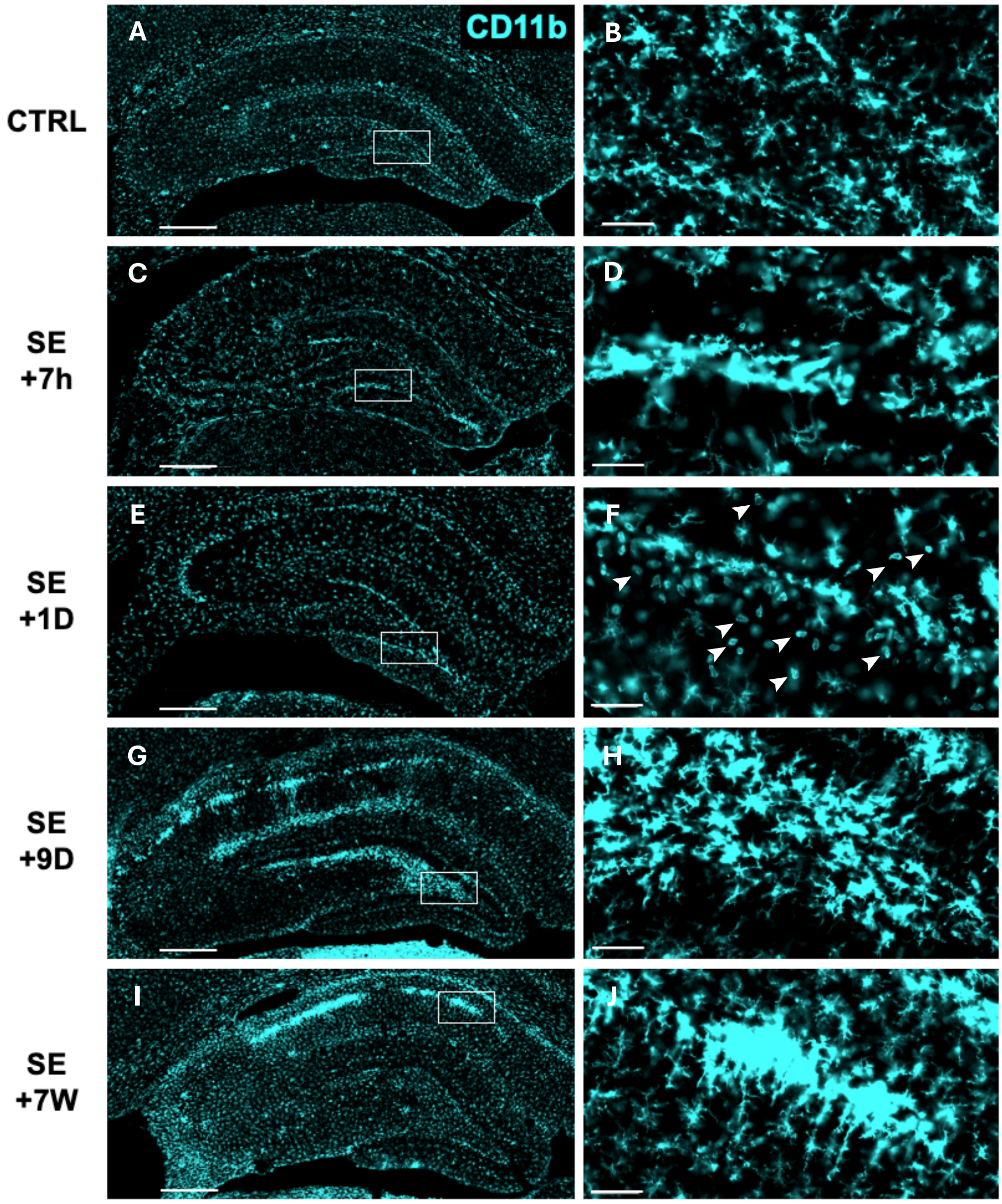

**Figure S1 – Evolution of microglial cell activation in the hippocampus after pilocarpine induced-SE.** CD11b (CBL1512Z, Millipore) was immunodetected for the visualization of microglia/monocytes in the hippocampus (Bregma -3.8mm) in control rats (**A-B**) and 7 hours (7h) (**C-D**), 1 day (1D) (**E-F**), 9D (**G-H**) and 7 weeks (7W) (**I-J**) post-SE. No differences were observed between the sections from the brain of control samples collected at P42 and those collected 7 weeks later. To simplify the figure, only images of the control samples collected at P42 are shown. Arrows in **F** point to round-shaped CD11b+ cells. Images were acquired with a slide scanner, objective ×20. Each image of the total hippocampus was zoomed in, with the zoomed area displayed in the adjacent image for a better visualization of cell morphology. Scale bars: left column, 500 µm; right column, 50µm.

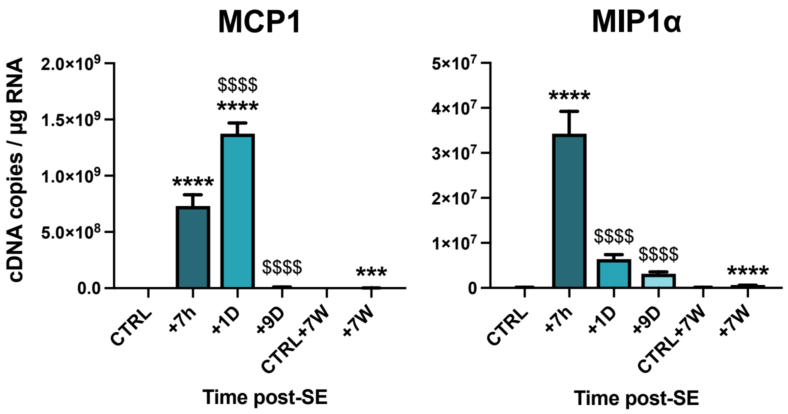

**Figure S2 – Transcript levels of chemokines are induced early during epileptogenesis in the hippocampal tissue.** MCP1 (monocyte attractant chemokine) and MIP1α (macrophage inflammatory protein α) mRNA levels were quantified using calibrated RT-qPCR in hippocampi microdissected from rat brains following SE (CTRL, n=6; SE+7h, n=6; SE+1D, n=6, SE+9D, n=7, CTRL+7W, n=6, SE+7W, n=8). Transcript levels are expressed as cDNA copies per µg of total RNA. Data measured 7h, 1 day (1D) and 9D post-SE were compared to P42 healthy controls, and those measured 7 weeks post-SE were compared to another age-matched control group. *: vs. respective CTRL; $: vs.SE+7h. */$, p<0.05; **/$$, p<0.01; ***/$$$, p<0.001; ****/$$$$, p<0.0001.

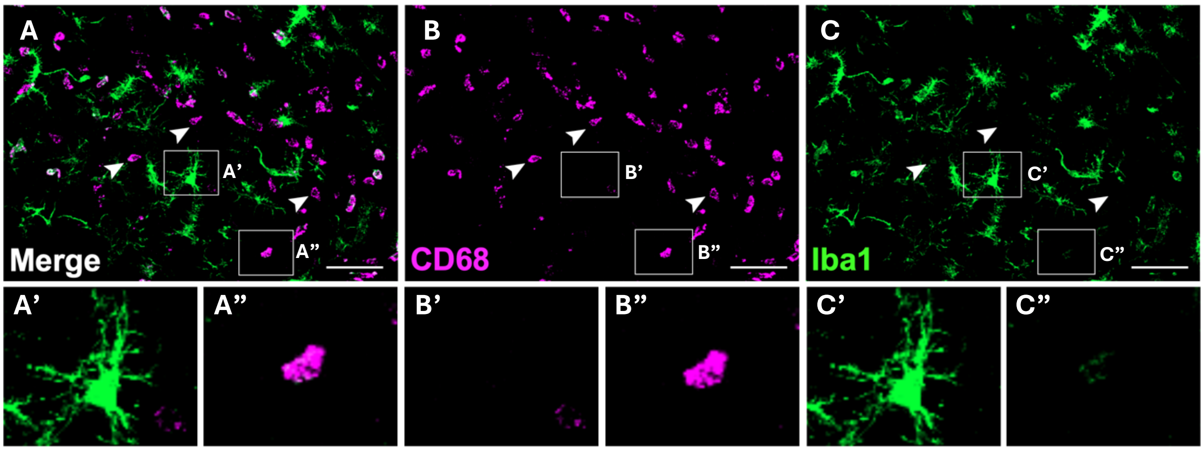

**Figure S3 – CD68 allows for detection of infiltrating monocytes in rat hippocampus following SE.** CD68 (magenta, MCA341GA, Bio-rad) and Iba1 (green, ab5076, abcam) were immunodetected in the dentate gyrus, 1 day post-SE. Arrows point out the presence of CD68-positive and Iba1-negative round cells, i.e., monocytes. An $Iba1^{+}CD68^{-}$ microglia-like cell is shown under each panel on the left (‘), and an $Iba1^{-}CD68^{+}$ monocyte-like cell is shwon under each panel on the right (“). Images were acquired with a confocal microscope, objective ×20. Scale bars: 50µm.

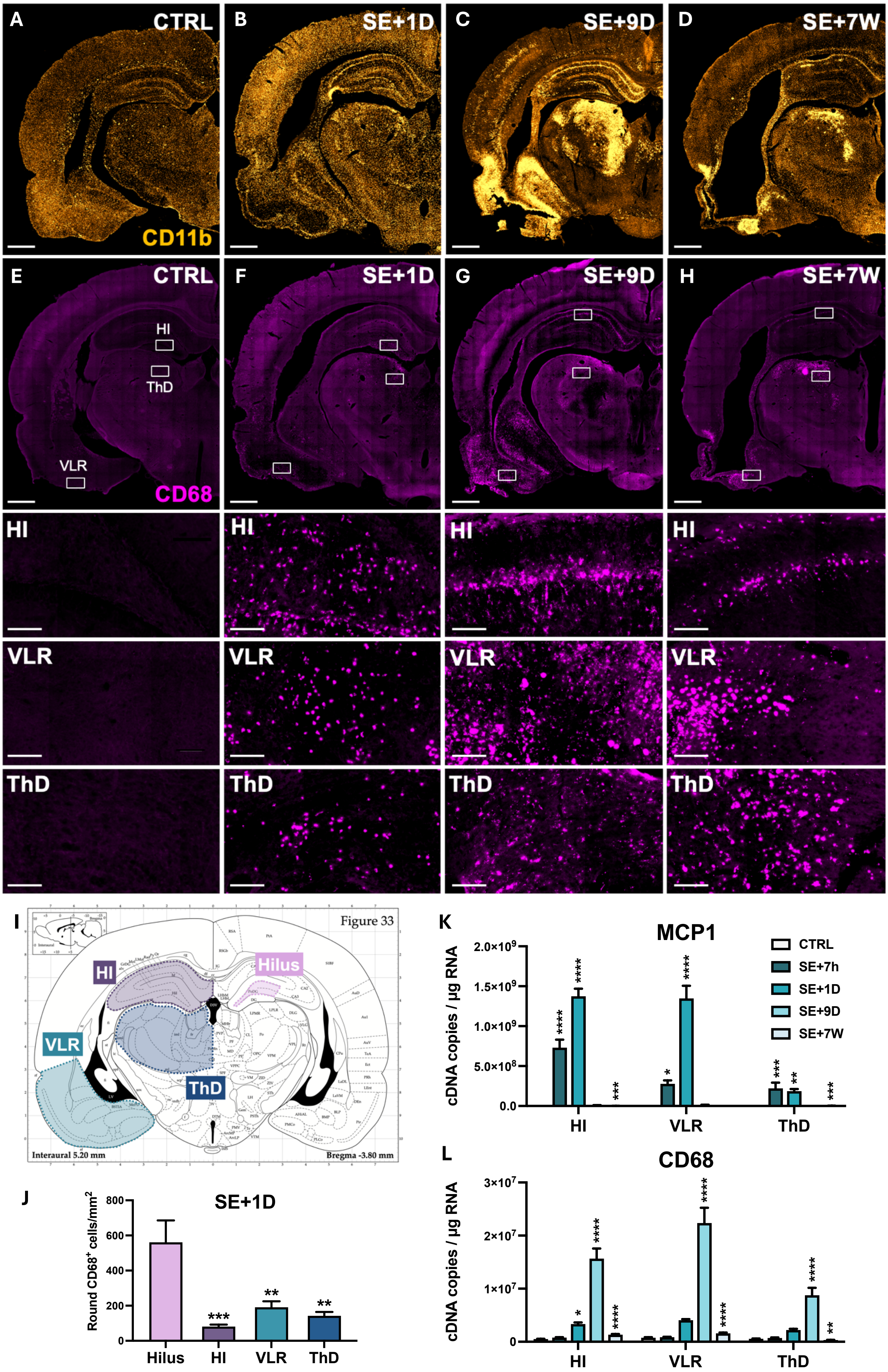

A CTRL
B SE+1D
C SE+9D
D SE+7W
CD11b
E CTRL
F SE+1D
G SE+9D
H SE+7W
HI
ThD
VLR
CD68
HI
VLR
ThD
I
Figure 33
HI
Hilus
VLR
ThD
Interaural 5.20 mm
Bregma -3.80 mm
J
SE+1D
Round CD68+ cells/mm2
Hilus
HI
VLR
ThD
K
MCP1
cDNA copies / µg RNA
CTRL
SE+7h
SE+1D
SE+9D
SE+7W
HI
VLR
ThD
L
CD68
cDNA copies / µg RNA
HI
VLR
ThD

**Figure S4 – Monocyte infiltration is strong in the hippocampus, ventral limbic region and dorsal thalamus and contributes to the formation of the microglial scar. A-H.** CD11b (CBL1512Z, Millipore) and CD68 (MCA341GA, Bio-rad) were immunodetected for visualization of myeloid cells and monocytes respectively, in whole brain slices (Bregma -3.8mm) in control rats (**A, E**) and 1D (**B, F**), 9D (**C, G**) and 7W (**D, H**) post-SE. CD11b labeling allows for the detection of the "microglial scar" from 9 days post-SE. CD68-positive monocytes are detectable up to 7 weeks post-SE in both the hippocampus ("HI"), the VLR, and the ThD, as shown in the magnifications. Images were acquired with a slide scanner, objective ×20. Scale bars: 1000 µm for whole brain images, 100 µm for magnifications. **I-J**. Round CD68-positive cells were automatically counted in the hilus, the total hippocampus, the VLR and the ThD 24h post-SE. Quantifications were made bilaterally and averaged for each animal. *: vs. Hilus. **K-L**. MCP1 (monocyte attractant chemokine) and CD68 (monocyte marker) mRNA levels were quantified using calibrated RT-qPCR in both hippocampi, ventral limbic region (VLR) comprising the piriform cortex, the amygdala and the insular agranular cortex, and the dorsal thalamus (ThD) microdissected from rat brains following SE (CTRL, n=6; SE+7h, n=6; SE+1D, n=6, SE+9D, n=7, CTRL+7W, n=6, SE+7W, n=8). Transcript levels are expressed as cDNA copies per µg of total RNAs. Data measured at 7h, 24h and 9 days post-SE were compared to P42 healthy controls, and those measured 7 weeks post-SE were compared to another age-matched control group. Only statistic comparisons with respective controls are presented. Details of statistical tests, including inter-region comparisons, are presented in Table S2. All data are presented as mean + SEM. *: vs. respective CTRL. *, $p<0.05$; **, $p<0.01$; ***, $p<0.001$; ****, $p<0.0001$

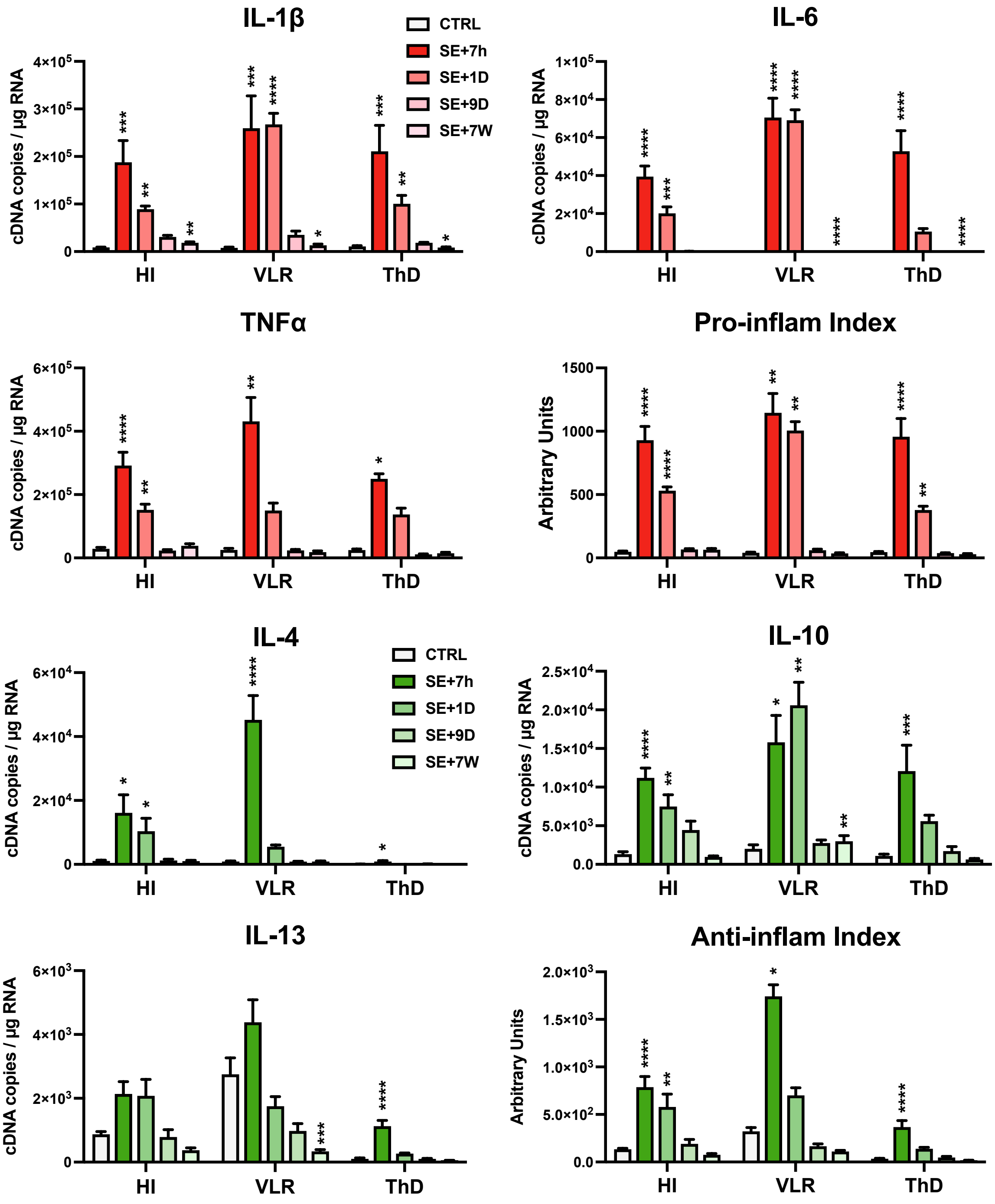

**Figure S5 – Transcriptional response of inflammatory markers in limbic brain regions during epileptogenesis and chronic epilepsy following SE induced by pilocarpine at P42 in rats.** mRNA levels of prototypical cytokine and cellular markers were quantified using calibrated RT-qPCR in the hippocampus, ventral limbic region (VLR) and dorsal thalamus (ThD) microdissected from rat brains following SE (CTRL, n=6; SE+7h, n=6; SE+1D, n=6, SE+9D, n=7, CTRL+7W, n=6, SE+7W, n=8). Transcript levels are expressed as cDNA copies per µg of total RNA. Pro- and anti-inflammatory indexes, calculated as described in the Method section, were based on IL-1β, IL-6 and TNFα (pro-inflammatory) and IL-4, IL-10 and IL-13 (anti-inflammatory) mRNA levels, expressed in arbitrary units. Data at 7h, 1 day and 9 days post-SE were compared to P42 healthy controls, while data at 7 weeks post-SE were compared to age-matched controls. Only statistical comparisons with respective controls are shown. All other comparisons, including inter-region comparisons, are presented in Table S2. Data are presented as mean + SEM. *: vs. respective CTRL.

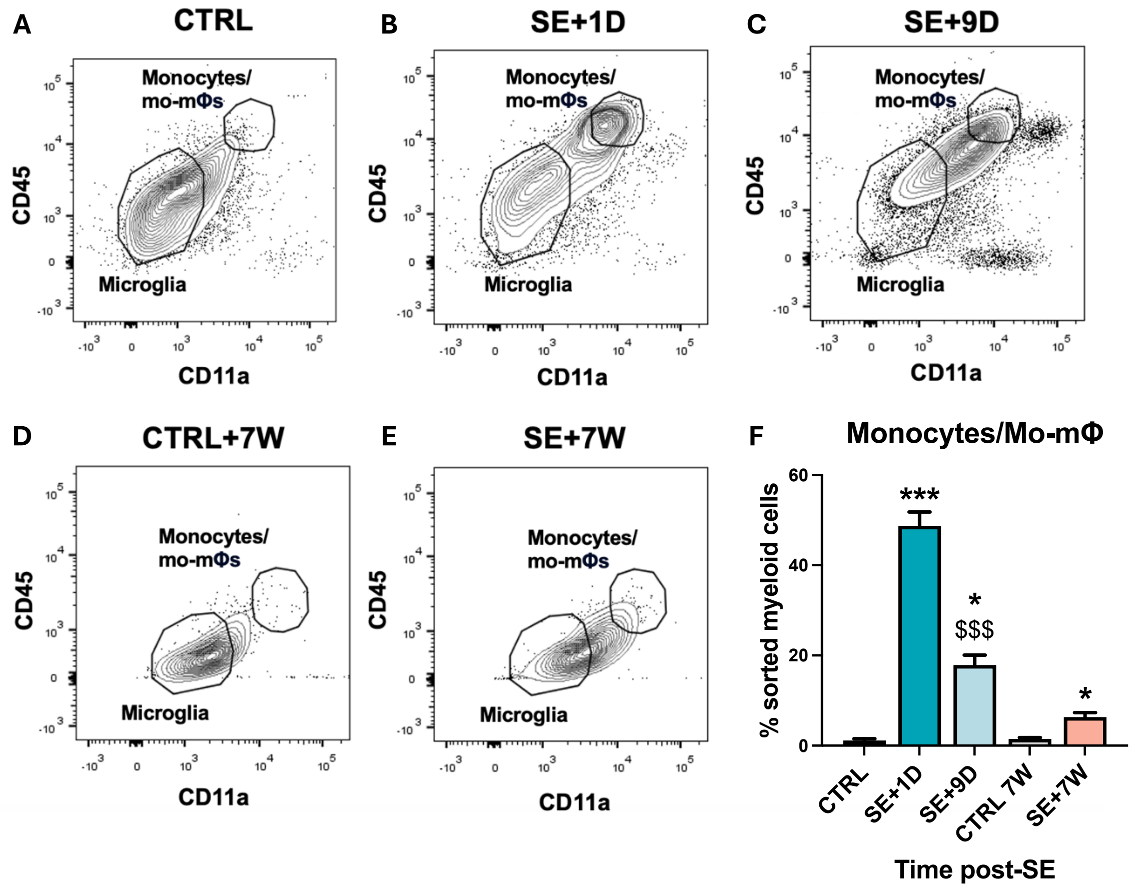

**Figure S6 – FACS-based sorting of microglia and monocytes/mo-mΦs up to 7 weeks post-SE. A-E.** Microglia ($CD11b^{+}CD45^{lo}CD11a^{lo}$) and monocytes/mo-mΦs ($CD11b^{+}CD45^{hi}CD11a^{hi}$) were sorted by FACS after CD11b MACS enrichment from the hippocampus, dorsal thalamus and ventral limbic region of rats (CTRL, n=2; SE+1D, n=3; SE+9D, n=3; CTRL 7W, n=3; SE+7W, n=3). **F**. The percentage of $CD11b^{+}CD45^{hi}CD11a^{hi}$ cells among all sorted cells ($CD11b^{+}CD45^{hi}CD11a^{hi}$ + $CD11b^{+}CD45^{lo}CD11a^{lo}$) was quantified. Data at 7h, 1 day (1D) and 9D post-SE were compared to P42 healthy controls ("CTRL"), while data at 7 weeks post-SE were compared to age-matched controls ("CTRL+7W"). Details of statistical tests are provided in supplementary table S2. Results are presented as mean + SEM. *: vs. respective CTRL; $: vs.SE+1D. */$, $p<0.05$; **/$$, $p<0.01$; ***/$$$, $p<0.001$; ****/$$$$, $p<0.0001$.

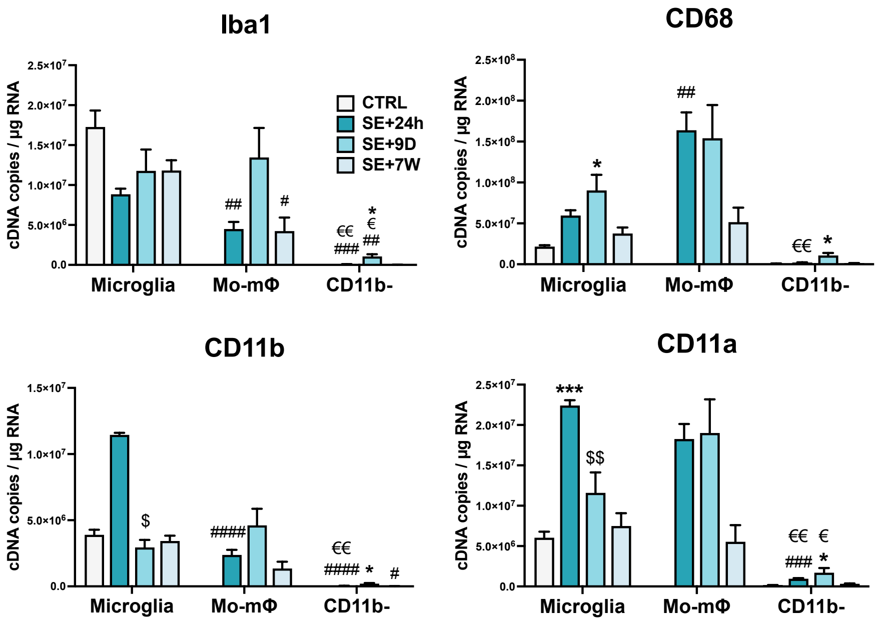

**Figure S7 – Transcript levels of cellular markers in sorted microglia, monocytes/mo-mΦ and CD11b-negative cells after SE.** CD11b- cells, microglia and monocytes/mo-mΦ were sorted using MACS and FACS. Transcript levels of cellular markers (Iba1, CD68, CD11b and CD11a) were quantified by calibrated RT-qPCR in these sorted cell populations (CTRL, n=2; SE+1D, n=3; SE+9D, n=3; CTRL 7W, n=3; SE+7W, n=3). Data at 7h, 1 day (1D) and 9D post-SE were compared to P42 healthy controls ("CTRL"), while data at 7 weeks post-SE were compared to age-matched controls ("CTRL+7W"). Statistical details are available in Table S2. Results are shown as mean + SEM. "Time post-SE" factor: *, vs. respective CTRL; $, vs. respective SE+1D; "cell type" factor: #, vs. respective microglia; €, vs. respective mo-mΦ. 1 symbol, $p<0.05$; 2 symbols, $p<0.01$; 3 symbols, $p<0.001$; 4 symbols, $p<0.0001$.

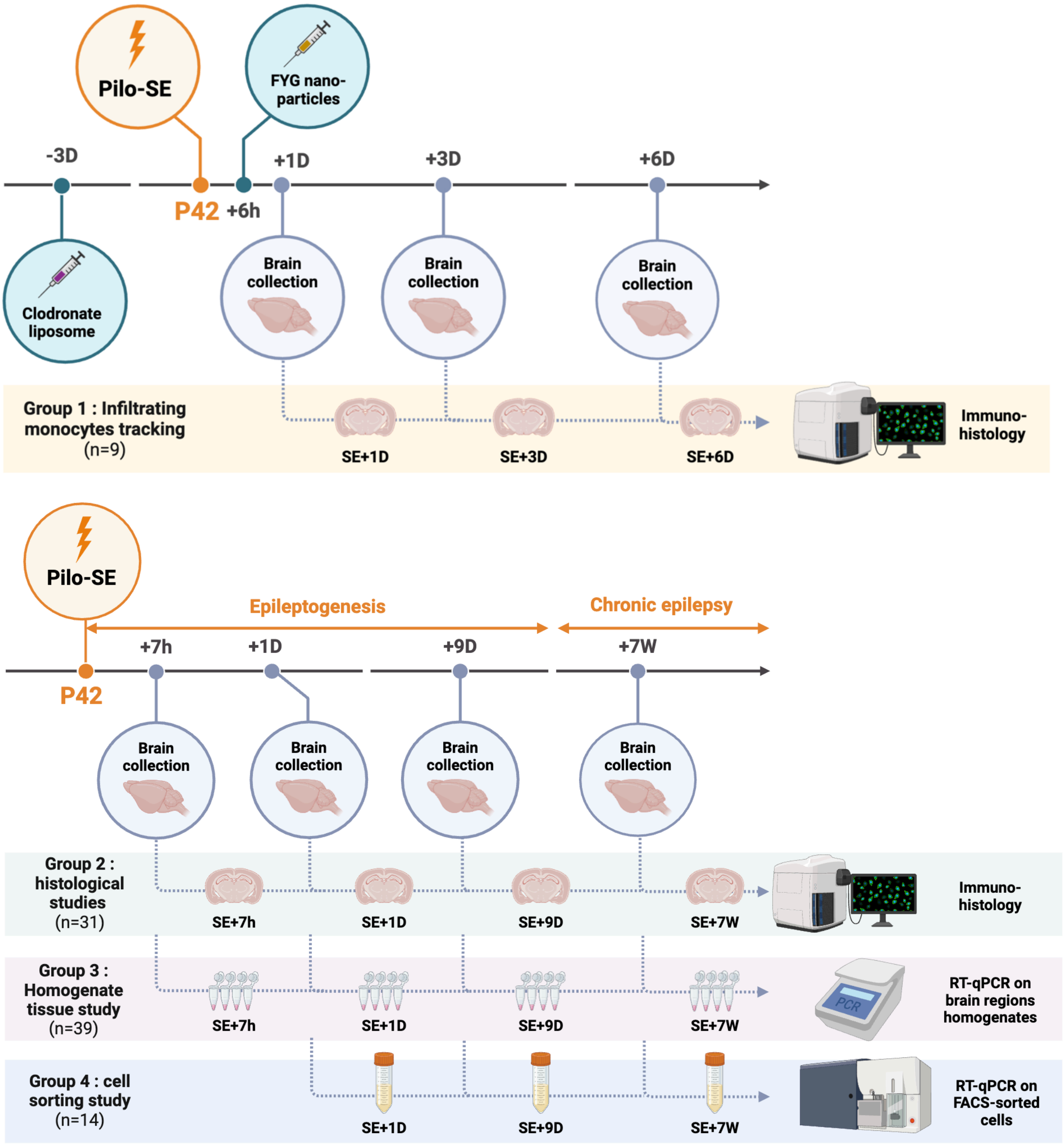

**Figure S8 – Experimental design.** Four groups of rats were subjected to pilocarpine-induced status epilepticus (SE) at P42. The results obtained during the epileptogenesis phase, between 7 hours and 9 days post-SE were compared with those of control rats at P42. Similarly, the results from the chronic phase of epilepsy, 7 weeks post-SE, were compared with those of age-matched controls (P42+7 weeks).

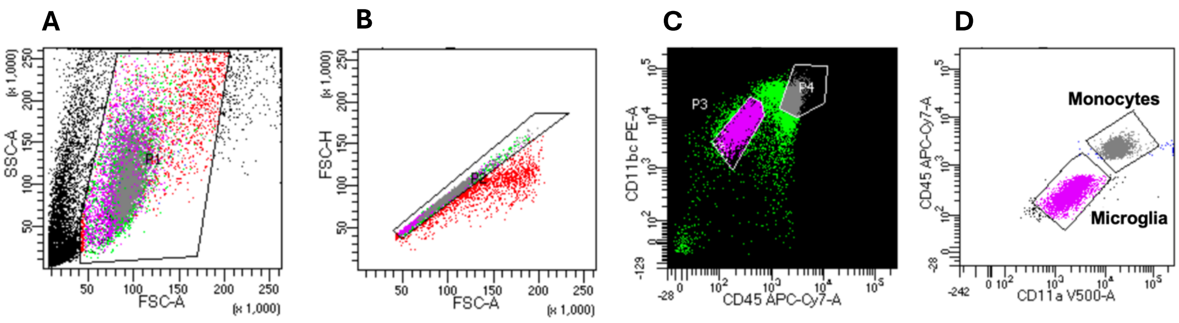

**Figure S9 – Gating strategy for sorting microglia and monocyte/ mo-mΦs sorting.** FACS was used to sort microglia (CD11b$^{+}$CD45$^{lo}$CD11a$^{lo}$) and monocytes/mo-mΦs (CD11b$^{+}$CD45$^{hi}$CD11a$^{hi}$) from a CD11b-positive cell-enriched population. **A**. Live cells were gated based on size and shape. **B**. Singlets were selected using the diagonal of the FSC-A /FSC-H plot. **C**. CD45 expression distinguished microglia (CD45$^{lo}$) from monocytes/mo-mΦs (CD45$^{hi}$) within CD11b-positve cells. **D**. CD11a expression further separated microglia (CD45$^{lo}$CD11a$^{lo}$, magenta) from monocytes/mo-mΦs (CD45$^{hi}$CD11a$^{hi}$, grey). Gates were defined using a SE+1D sample (example shown) and applied consistently across all samples.

### Method details

#### Brain collections

All rats were deeply anesthetized with isoflurane (4%) and then administered a lethal dose of pentobarbital (100 mg/kg, i.p.; Euthasol). **For histological studies (Groups 1 and 2)**, animals were intracardially perfused at a flow rate of 30mL/min, first with ice-cold saline to clear blood from brain vessels, followed by 4% paraformaldehyde in 0.1 M phosphate buffer (PB) to fix tissue. **For RT-qPCR analysis (Group 3)**, animals were intracardially perfused with ice-cold saline (30mL/min) to remove blood from brain vessels. The hippocampus, ventral limbic region and dorsal thalamus were rapidly microdissected on ice, frozen in liquid nitrogen, and stored at -80°C. **For flow cytometry analysis (Group 4)**, transcardiac perfusion was performed using ice-cold Dulbecco's PB saline (DPBS) supplemented with a cocktail of transcription and translation inhibitors (so-called inhibition cocktail), including Actinomycin D (3µM), Anisomycin (100 µM) and Triptolide (10 µM). Once perfused, brain tissues were quickly dissected and placed in the same inhibitor-enriched buffer until all perfusions were completed.

#### Immunohistology

**Tissue processing** – 40 µm-thick coronal sections were cut from frozen brains using a cryomicrotome (Leica CM1850; -22°C). Sections were collected in 0.1M Phosphate Buffer Saline, (PBS) and transferred into a cryopreservative solution composed of 19.5 mM $NaH_2PO_4 \bullet 2H_2O$, 19.2 mM NaOH, 30% glycerol and 30% ethylene glycol, and then stored at -20°C. Before starting any labeling protocol, free-floating sections were rinsed 3 times with PBS.

**Fluorescent immunolabelling** – Following permeabilization and saturation with 0.3% Triton 100X and 3% Donkey normal serum, free-floating sections were incubated with primary antibodies. Detection and morphology appreciation of myeloid cells was performed with mouse anti-Cluster of differentiation 11b (CD11b) antibody (1:2000, CBL1512Z, Millipore). Detection of microglia was performed with goat anti-Ionized calcium binding adaptor molecule 1 (Iba1) antibody (1:500, ab5076, Abcam). Infiltrating monocytes and monocyte-macrophages (mo-MΦ) were labelled with mouse anti-CD68 antibody (1:1000, MCA341GA, Bio-rad). Alexa-Fluor-488-conjugated donkey anti-mouse antibody was used for CD11b labelling (1:1000, A-21202, Molecular Probes). For fluorescent dual immunolabeling of Iba-1 and CD68, sections were incubated with an Alexa-Fluor-488-conjugated donkey anti-goat antibody (1:1000, A-11055, Molecular Probes) and an Alexa-Fluor-647-conjugated donkey anti-mouse antibody (1:1000, A-31571, Molecular Probes). Nuclei were stained with DAPI (300 nM, Molecular Probes). Sections were mounted on SuperFrost Plus slides and coverglassed with Prolong Diamond Antifade reagent (Molecular Probes)

**Microscopy** – Whole sections were scanned with a Carl Zeiss Axio Scan.Z1 Digital Slide Scanner (ZEISS) with a ×20 lens on a 6 µm stack, using the pilot Zen (ZEISS), or observed with a TCS SP5X confocal microscopy system (Leica). Images were processed on with Fiji software (ImageJ).

**Molecular biology**

**RNA extraction –** Tissues (Group 2) were crushed in 250 µL of molecular biology grade water (Eurobio) using Tissue-Lyser II (Qiagen) according to the manufacturer's instructions. Total RNAs from brain structures were extracted using Tri-Reagent LS (Molecular Research Center, #TS120). Genomic DNA was removed using Turbo-DNA-free (Ambion, #1907) and total RNAs were purified with RNeasy mini kit (Qiagen, #74104). For sorted cells (Group 3), total RNAs were extracted and purified using the RNeasy Plus Micro Kit (Qiagen, #74034) according to manufacturer's instruction. RNA concentration was determined for each sample on the BioDrop® µLite.

**Reverse transcription and real-time quantitative PCR –** Total tissue RNAs were reverse transcribed to complementary DNA (cDNA) in a total volume of 10 µL using a mix of oligo dT and random primers as well as PrimeScript RT Reagent Kit (Takara, #RR037A) according to manufacturer's instructions. In RT reaction, 300 000 copies of a synthetic external non-homologous poly(A) standard messenger RNA (SmRNA; A. Morales and L. Bezin, patent WO2004.092414) were added to normalize the RT step, as previously described (*43*). cDNA was diluted 1:13 with nuclease free Eurobio water and stored at -20°C until further use. Each cDNA of interest was amplified from 5 µL of the diluted RT reaction by real-time PCR, using the Rotor-Gene Q thermocycler (Qiagen), the SYBR Green PCR kit (Qiagen, #208052) and oligonucleotide primers (Eurogentec) specific to the targeted cDNA. The sequences of the specific forward and reverse primer pairs were constructed using Primer-BLAST (NCBI). Primers used are listed in Table S1. cDNA copy number detected was determined using a calibration curve, and results were expressed as cDNA copy number/µg tot RNA.

The pro-inflammatory index (PI-I) and anti-inflammatory index (AI-I) were calculated based on specific sets of inflammatory genes: IL-1β, IL-6, TNFα for PI-I, and IL-4, IL-10, IL-13 for AI-I. For each sample, the expression level of each transcript (in cDNA copies) was normalized as a percentage of the average copy number across all samples. This normalization allows the integration of transcripts with vastly different expression levels within each index, ensuring that highly expressed transcripts do not overshadow variations in those expressed at lower levels. Each index was then computed by summing the percentages of the transcripts contributing to its composition, with the final results expressed in arbitrary units (A.U.), according to the following formula:

$$PI = \sum_{i=a}^{n} \frac{cDNA\ copy\ nbr\ for\ transcript\ i}{average\ cDNA\ copy\ nbr\ for\ transcript\ i}$$

with i = IL-1β or IL-6 or TNFα for PI-I and i = IL-4 or IL-10 or IL-13 for AI-I.

**Flow cytometry**

Samples were processed for tissue dissociation and cell sorting immediately following brain collection as quickly as possible. To prevent any artifactual *ex vivo* gene expression changes during brain dissociation and cell sorting procedures, all buffers and solutions used during the process (from animal perfusion to sorted cell flash freezing) were supplemented with a cocktail composed of Actinomycin D (3 µM, Tocris #1229/10), Anisomycin (100 µM, Tocris #1290/50) and Triptolide (10 µM, Tocris #3253/10) i.e. transcription and translation inhibitors (*24*). All steps were performed on ice or using pre-chilled refrigerated centrifuge set to 4°C with all buffers/solutions pre-chilled before addition to samples to further limit cell activation. Buffer 1 (B1) was composed of DPBS (Thermofisher #14040-117) and inhibitor cocktail. Buffer 2 (B2) was composed of DPBS, BSA 0,5% (Sigma #A2153) and inhibitor cocktail.

**Brain tissue dissociation** – Once collected, tissues were cut in smaller pieces with a scalpel and processed for dissociation using Miltenyi's Adult Brain Dissociation Kit (#130-107-677) according to manufacturer's instruction. Inhibitor cocktail was added in each reagent. Briefly, samples were added to gentleMACS C Tubes (Miltenyi #130-093-237) with the enzyme mixes and placed in gentleMACS OctoDissociatior with heaters (Miltenyi #130-096-427), running program 37C_ABDK_01. Once program finished, samples were briefly spun before being filtered through 70 µm cell strainer

(Thermofisher #11597522). Samples were washed with B1 and spun to pellet cells. To clear cell solution, cell pellets were resuspended and overlaid with appropriate volume of Miltenyi Debris Removal Solution according to manufacturer's protocol. Debris were removed from top layer and solution was diluted with B1 and spun to pellet cells. Cells collected were resuspended in B1 and counted manually (with trypan blue) before magnetic sorting.

**CD11b-positive cells magnetic enrichment** – To increase Fluorescence-Activated Cell Sorting (FACS) yields and efficiency, cell suspensions were first enriched using the Magnetic-Activated Cell Sorting (MACS) technique, magnetically separating CD11b-positive cells (microglia and infiltrating monocytes) for subsequent FACS, from CD11b-negative cells (remaining brain cells, i.e. neurons, astrocytes, oligodendrocytes, endothelial cells, etc.) for direct freezing. In the following steps, cells were suspended in B2. Fc receptors were blocked with anti-CD32 antibody (BD Biosciences #550271). CD11b-positive cells were then enriched using CD11b/c MicroBeads according to manufacturer's instructions (Miltenyi #130-105-634). Briefly, brain cells were incubated with CD11b/c MicroBeads and applied onto MS columns (Miltenyi #130-042-201) in the magnetic field of OctoMACS Separator (Miltenyi #130-042-109). CD11b-positive fractions were collected and subjected to the FACS protocol. Unlabelled cells were spun and dry cell pellets were flash frozen and stored at -80°C for further analysis.

**FACS** – Cells were incubated in B2 with anti-CD11b/c PE-conjugated (1:50, BD Biosciences #554862), anti-CD11a BV510-conjugated (1:50, BD Biosciences #744999) and anti-CD45 APCCy7-conjugated (1:50, Biolegend #202216) antibodies. Microglia ($CD11b^{+}CD45^{lo}CD11a^{lo}$) and infiltrating monocytes ($CD11b^{+}CD45^{hi}CD11a^{hi}$) were sorted with BD FACS Aria™ III Cell Sorter (BD Biosciences), and acquired with the BD FACSDiva 9.4 software. Further analyses were made with FlowJo software. Gating strategy on CD11-positive cells is shown in **Figure S9**.

**Statistical Analysis**

Statistical analyses were performed using Prism 10.0 software (GraphPad, USA). Results are presented as mean + SEM (standard error of the mean). Differences with a p-value<0.05 ($p<0.05$) were considered to be statistically significant. The Shapiro–Wilk test and quantile–quantile plot were used to assess normal distribution of the data. For the data with normal distribution, the statistical significance was assessed by t-test, one-way or two-way ANOVA analysis, followed with Tukey's post-hoc test for multiple comparisons. For non-normal distribution data, the statistical significance was assessed by Kruskal-Wallis' test, followed with Dunn's post-hoc test for multiple comparisons or Mann-Whitney test for two groups comparisons. Simple linear regression was used to assess the association between two variables. Details of statistic tests for each figure are presented in **Table S2**.

**Table S1 – Sequences of primer pairs used for qPCR** (*Rattus Norvegicus*)

| Target cDNA | Forward primer sequence | Reverse primer sequence | GenBank Reference |
|---|---|---|---|
| **Arg1** | 5’ TCC AAG CCA AAG CCC ATA GAG 3’ | 3’ CTT TGT ATG TTA CAC TCT CTG 5’ | NM_017134.3 |
| **CD11a** | 5’ GAA GGA TCA CCA AGG GAC CAA 3’ | 3’AGG GGA TCT CCT TAG TGA CTG 5’ | NM_001033998.2 |
| **CD11b** | 5’ ACT CTG ATG CCT CCC TTG G 3’ | 3’ TCC TGG ACA CGT TGT TCT CA 5’ | NM_012711.1 |
| **CD68** | 5’ CTT TCT CCA GCA ATT CAC CTG 3’ | 3’ ACT GGC GCA AGA GAA GCA 5’ | NM_001031638.1 |
| **CD206** | 5’ TTT GAG TGG AGC GAT GGG AC 3’ | 3’ CAT GCC GTT TCC AGC CTT TC 5’ | NM_001106123.2 |
| **Iba1** | 5’ CCA GCG TCT GAG CTA TG 3’ | 3’ CCA GCA TTC GCT TCA AGG AC 5’ | NM_01716.3 |
| **IL-10** | 5’ TTG AAC CAC CCG GCA TCT AC 3’ | 3’ CCA AGG AGT TGC TCC CGT TA 5’ | NM_012854.2 |
| **IL-13** | 5’ AGT CCT GGC TCT CGC TTG 3’ | 3’ GAT GTG GAT CTC CGC ACT G 5’ | NM_053828.1 |
| **IL-1β** | 5’ TGT GAT GAA AGA CGG CAC AC 3’ | 3’ CTT CTT CTT TGG GTA TTG TTT GG 5’ | NM_031512.2 |
| **IL-4** | 5’ GTA GAG GTG TCA GCG GTC TG 3’ | 3’ TTC AGT GTT GTG AGC GTG GA 5’ | NM_201270.1 |
| **IL-6** | 5’ CCC TTC AGG AAC AGC TAT GAA 3’ | 3’ ACA ACA TCA GTC CCA AGA AGG 5’ | NM_012589.1 |
| **MCP1** | 5’ CGG CTG GAG AAC TAC AAG AGA 3’ | 3’ TCT CTT GAG CTT GGT GAC AAA TA 5’ | NM_57441.1 |
| **MIP1α** | 5’ TCC ACG AAA ATT CATT GCT G 3’ | 3’ AGA TCT GCC GGT TTC TCT TG 5’ | NM_013025.2 |
| **TNFα** | 5’ TGA ACT TCG GGG TGA TCG 3’ | 3’ GGG CTT GTC ACT CGA GTT TT 5’ | NM_012675.3 |
| **SmRNA** | 5’ CGG GAC AAG AAG GTG GAA G 3’ | 5’ AGT CTG CAG TGA GTC GAA GAA A 3’ | WO2004.092414 |

**Abbreviations**: Arg1, arginase 1; CD, cluster of differentiation; Iba1, ionized calcium-binding adapter molecule 1; IL, interleukin; MCP1, monocyte chemoattractant protein 1; MIP1α, macrophage inflammatory protein α; TNFα, tumor necrosis factor α; SmRNA, standard messenger RNA.

**Table S2 – Details of statistic tests.** Normality of data distribution was tested using Shapiro–Wilk test and quantile–quantile plots (“Normality?” column) and the statistic test was chosen accordingly for each group.

| Figure 1 | | | | | | | Normality ? | Test | Post-hoc |
|---|---|---|---|---|---|---|---|---|---|
| | | | p-value | | | | | | |
| | | | vs. CTRL (*) | | vs. SE+1D ($) | | | | |
| 1J | CD11b | SE+7h | 0.9967 | ns | | | Yes | One-way ANOVA | Tukey's |
| | | SE+1D | <0.0001 | **** | <0.0001 | $$$$ | | | |
| | | SE+6D | 0.9964 | ns | >0.9999 | ns | | | |
| 1N | CD68 | SE+7h | >0.9999 | ns | | | Yes | One-way ANOVA | Tukey's |
| | | SE+1D | <0.0001 | **** | 0.0002 | $$$ | | | |
| | | SE+6D | 0.6620 | ns | 0.7495 | ns | | | |

| Figure 2 | | | | | | Normality ? | Test | Post-hoc |
|---|---|---|---|---|---|---|---|---|
| | | | | p-value | | | | |
| 2K | Hilus | Round vs. ramified | CTRL | >0.9999 | ns | Yes | Two-way RM ANOVA | Tukey’s |
| | | | +7h | 0.9482 | ns | | | |
| | | | +1D | <0.0001 | **** | | | |
| | | | +9D | <0.0001 | **** | | | |
| | | | +7W | 0.0531 | ns | | | |
| | | Round cells | CTRL vs. SE+7h | >0.9999 | ns | | | |
| | | | CTRL vs. SE+1D | <0.0001 | **** | | | |
| | | | CTRL vs. SE+9D | 0.0998 | ns | | | |
| | | | CTRL vs. SE+7W | 0.9964 | ns | | | |
| | | Ramified cells | CTRL vs. SE+7h | >0.9999 | ns | | | |
| | | | CTRL vs. SE+1D | >0.9999 | ns | | | |
| | | | CTRL vs. SE+9D | <0.0001 | **** | | | |
| | | | CTRL vs. SE+7W | 0.1927 | ns | | | |
| 2L | CA1 | Round vs. ramified | CTRL | >0.9999 | ns | Yes | Two-way RM ANOVA | Tukey’s |
| | | | +7h | 0.9475 | ns | | | |
| | | | +1D | 0.8051 | ns | | | |
| | | | +9D | <0.0001 | **** | | | |
| | | | +7W | 0.2925 | ns | | | |
| | | Round cells | CTRL vs. SE+7h | >0.9999 | ns | | | |
| | | | CTRL vs. SE+1D | 0.9996 | ns | | | |
| | | | CTRL vs. SE+9D | 0.8871 | ns | | | |
| | | | CTRL vs. SE+7W | 0.9965 | ns | | | |
| | | Ramified cells | CTRL vs. SE+7h | >0.9999 | ns | | | |
| | | | CTRL vs. SE+1D | >0.9999 | ns | | | |
| | | | CTRL vs. SE+9D | >0.0001 | **** | | | |
| | | | CTRL vs. SE+7W | 0.7371 | ns | | | |

| Figure 3 | | | | | | | | | |
|---|---|---|---|---|---|---|---|---|---|
| | | | p-value | | | | Normality ? | Test | Post-hoc |
| | | | vs. CTRL (*) | | vs. SE+7h ($) | | | | |
| 3A | IL-1β | SE+7h | <0.0001 | **** | | | Yes | One-way ANOVA | Tukey's |
| | | SE+1D | 0.0009 | *** | 0.2030 | ns | | | |
| | | SE+9D | 0.9207 | ns | <0.0001 | $$$$ | | | |
| | | SE+7W | 0.0016 | ** | | | Yes | Unpaired t-test | |
| 3B | IL-6 | SE+7h | <0.0001 | **** | | | Yes | One-way ANOVA | Tukey's |
| | | SE+1D | <0.0001 | **** | <0.0001 | $$$$ | | | |
| | | SE+9D | >0.9999 | ns | <0.0001 | $$$$ | | | |
| | | SE+7W | <0.0001 | **** | | | Yes | Unpaired t-test | |
| 3C | TNFα | SE+7h | 0.086 | * | | | No | Krukall Wallis | Dunn's |
| | | SE+1D | 0.4751 | ns | >0.9999 | ns | | | |
| | | SE+9D | >0.9999 | ns | 0.0002 | $$$ | | | |
| | | SE+7W | 0.9497 | ns | | | No | Mann-Whitney | |
| 3D | Pro-inflam index | SE+7h | <0.0001 | **** | | | Yes | One-way ANOVA | Tukey's |
| | | SE+1D | <0.0001 | **** | 0.0004 | $$$ | | | |
| | | SE+9D | 0.9986 | ns | <0.0001 | $$$$ | | | |
| | | SE+7W | 0.2038 | ns | | | Yes | Unpaired t-test | |
| 3E | IL-4 | SE+7h | 0.0050 | ** | | | No | Krukall Wallis | Dunn's |
| | | SE+1D | 0.1485 | ns | >0.9999 | ns | | | |
| | | SE+9D | >0.9999 | ns | 0.0014 | $$ | | | |
| | | SE+7W | 0.7937 | ns | | | Yes | Unpaired t-test | |
| 3F | IL-10 | SE+7h | 0.0023 | ** | | | No | Krukall Wallis | Dunn's |
| | | SE+1D | 0.0017 | ** | >0.9999 | ns | | | |
| | | SE+9D | 0.9402 | ns | 0.1203 | ns | | | |
| | | SE+7W | 0.0426 | * | | | No | Mann-Whitney | |
| 3G | IL-13 | SE+7h | 0.0016 | ** | | | Yes | One-way ANOVA | Tukey's |
| | | SE+1D | 0.9730 | ns | 0.0041 | $$ | | | |
| | | SE+9D | 0.1426 | ns | <0.0001 | $$$$ | | | |
| | | SE+7W | 0.0030 | ** | | | Yes | Unpaired t-test | |
| 3H | Anti-inflam index | SE+7h | <0.0001 | **** | | | Yes | One-way ANOVA | Tukey's |
| | | SE+1D | <0.0001 | **** | <0.0001 | $$$$ | | | |
| | | SE+9D | 0.9226 | ns | <0.0001 | $$$$ | | | |
| | | SE+7W | 0.1044 | ns | | | Yes | Unpaired t-test | |

| Figure 4 | | | | | | | | |
|---|---|---|---|---|---|---|---|---|
| | | | | p-value | | Normality ? | Test | Post-hoc |
| 4B | IL-1β | Microglia | CTRL vs. SE+1D | 0.0169 | * | Yes | One-way ANOVA | Tukey's |
| | | | CTRL vs. SE+9D | 0.9684 | ns | | | |
| | | | SE+1D vs. SE+9D | 0.0130 | $ | | | |
| | | Mo-mΦ | SE+1D vs. SE+9D | 0.0022 | $$ | Yes | Unpaired t-test | |
| | | CD11b- | CTRL vs. SE+1D | 0.0417 | * | No | Krukall Wallis | Dunn's |
| | | | CTRL vs. SE+9D | 0.7907 | ns | | | |
| | | | SE+1D vs. SE+9D | 0.4008 | ns | | | |
| | | CTRL | Microglia vs. CD11b- | 0.2000 | ns | No | Mann-Whitney | |
| | | SE+1D | Microglia vs. Mo-mΦ | 0.2282 | ns | Yes | RM One-way ANOVA | Tukey's |
| | | | Microglia vs. CD11b- | 0.0192 | # | | | |
| | | | Mo-mΦ vs. CD11b- | 0.1036 | ns | | | |
| | | SE+9D | Microglia vs. Mo-mΦ | 0.2221 | ns | Yes | RM One-way ANOVA | Tukey's |
| | | | Microglia vs. CD11b- | 0.1928 | ns | | | |
| | | | Mo-mΦ vs. CD11b- | 0.9899 | ns | | | |
| | IL-1β 7W | CTRL 7W vs. SE+7W | Microglia | 0.0565 | ns | Yes | Unpaired t-test | |
| | | | CD11b- | 0.2842 | ns | Yes | Unpaired t-test | |
| | | CTRL 7W | Microglia vs. CD11b- | 0.0223 | # | Yes | Paired t-test | |
| | | SE+7W | Microglia vs Mo-mΦ | 0.8504 | ns | Yes | RM One-way ANOVA | Tukey's |
| | | | Microglia vs CD11b- | 0.1299 | ns | | | |
| | | | Mo-mΦ vs CD11b- | 0.0760 | ns | | | |
| 4C | IL-6 | Microglia | CTRL vs. SE+1D | 0.5391 | ns | No | Krukall Wallis | Dunn's |
| | | | CTRL vs. SE+9D | 0.5391 | ns | | | |
| | | | SE+1D vs. SE+9D | 0.0219 | $ | | | |
| | | Mo-mΦ | SE+1D vs. SE+9D | 0.0091 | $$ | Yes | Unpaired t-test | |
| | | CD11b- | CTRL vs. SE+1D | 0.1325 | ns | No | Krukall Wallis | Dunn's |
| | | | CTRL vs. SE+9D | >0.9999 | ns | | | |
| | | | SE+1D vs. SE+9D | 0.2003 | ns | | | |
| | | CTRL | Microglia vs. CD11b- | 0.2000 | ns | No | Mann-Whitney | |
| | | SE+1D | Microglia vs. Mo-mΦ | 0.3569 | ns | Yes | RM One-way ANOVA | Tukey's |
| | | | Microglia vs. CD11b- | 0.0978 | ns | | | |
| | | | Mo-mΦ vs. CD11b- | 0.0253 | € | | | |
| | | SE+9D | Microglia vs. Mo-mΦ | 0.3074 | ns | No | Friedman test | Dunn's |
| | | | Microglia vs. CD11b- | >0.9999 | ns | | | |
| | | | Mo-mΦ vs. CD11b- | >0.9999 | ns | | | |
| | IL-6 7W | CTRL 7W vs. SE+7W | Microglia | 0.2000 | ns | No | Mann-Whitney | |
| | | | CD11b- | 0.6041 | ns | Yes | Unpaired t-test | |
| | | CTRL 7W | Microglia vs. CD11b- | >0.9999 | ns | No | Wilcoxon | |
| | | SE+7W | Microglia vs Mo-mΦ | 0.6620 | ns | No | Friedman | Dunn's |
| | | | Microglia vs CD11b- | 0.6620 | ns | | | |
| | | | Mo-mΦ vs CD11b- | >0.9999 | ns | | | |
| 4D | TNFα | Microglia | CTRL vs. SE+1D | 0.0044 | ** | Yes | One-way ANOVA | Tukey's |
| | | | CTRL vs. SE+9D | 0.1536 | ns | | | |
| | | | SE+1D vs. SE+9D | 0.0007 | $$$ | | | |
| | | Mo-mΦ | SE+1D vs. SE+9D | 0.0515 | ns | Yes | Unpaired t-test | |
| | | CD11b- | CTRL vs. SE+1D | 0.3526 | ns | No | Krukall Wallis | Dunn's |
| | | | CTRL vs. SE+9D | >0.9999 | ns | | | |
| | | | SE+1D vs. SE+9D | >0.9999 | ns | | | |
| | | CTRL | Microglia vs. CD11b- | 0.2000 | ns | No | Mann-Whitney | |
| | | SE+1D | Microglia vs. Mo-mΦ | 0.0004 | ### | Yes | RM One-way ANOVA | Tukey's |
| | | | Microglia vs. CD11b- | 0.0003 | ### | | | |
| | | | Mo-mΦ vs. CD11b- | 0.8492 | ns | | | |
| | | SE+9D | Microglia vs. Mo-mΦ | 0.0582 | ns | Yes | RM One-way ANOVA | Tukey's |
| | | | Microglia vs. CD11b- | 0.0078 | ## | | | |
| | | | Mo-mΦ vs. CD11b- | 0.2574 | ns | | | |
| | TNFα 7W | CTRL 7W vs. SE+7W | Microglia | 0.0522 | ns | Yes | Unpaired t-test | |
| | | | CD11b- | 0.8595 | ns | Yes | Unpaired t-test | |

| | | | | | | | | |
|---|---|---|---|---|---|---|---|---|
| | | CTRL 7W | Microglia vs. CD11b- | <0.0001 | #### | Yes | Paired t-test | |
| | | SE+7W | Microglia vs Mo-mΦ | 0.0055 | ## | Yes | RM One-way ANOVA | Tukey's |
| | | | Microglia vs CD11b- | 0.0024 | ## | | | |
| | | | Mo-mΦ vs CD11b- | 0.6681 | ns | | | |
| 4E | MIP1α | Microglia | CTRL vs. SE+1D | 0.0003 | *** | Yes | One-way ANOVA | Dunn's |
| | | | CTRL vs. SE+9D | 0.7985 | ns | | | |
| | | | SE+1D vs. SE+9D | 0.0004 | $$$ | | | |
| | | Mo-mΦ | SE+1D vs. SE+9D | 0.0323 | $ | Yes | Unpaired t-test | |
| | | CD11b- | CTRL vs. SE+1D | 0.0626 | ns | No | Krukall Wallis | Dunn's |
| | | | CTRL vs. SE+9D | 0.6154 | ns | | | |
| | | | SE+1D vs. SE+9D | 0.73 | ns | | | |
| | | CTRL | Microglia vs. CD11b- | 0.2000 | ns | No | Mann-Whitney | |
| | | SE+1D | Microglia vs. Mo-mΦ | 0.0032 | ## | Yes | RM One-way ANOVA | Tukey's |
| | | | Microglia vs. CD11b- | 0.0015 | ## | | | |
| | | | Mo-mΦ vs. CD11b- | 0.2864 | ns | | | |
| | | SE+9D | Microglia vs. Mo-mΦ | 0.9966 | ns | Yes | RM One-way ANOVA | Tukey's |
| | | | Microglia vs. CD11b- | 0.0763 | ns | | | |
| | | | Mo-mΦ vs. CD11b- | 0.071 | ns | | | |
| | MIP1α 7W | CTRL 7W vs. SE+7W | Microglia | 0.0880 | ns | Yes | Unpaired t-test | |
| | | | CD11b- | 0.4111 | ns | Yes | Unpaired t-test | |
| | | CTRL 7W | Microglia vs. CD11b- | 0.0042 | ## | Yes | Paired t-test | |
| | | SE+7W | Microglia vs Mo-mΦ | 0.8385 | ns | Yes | RM One-way ANOVA | Tukey's |
| | | | Microglia vs CD11b- | 0.0937 | ns | | | |
| | | | Mo-mΦ vs CD11b- | 0.1668 | ns | | | |
| 4F | MCP1 | Microglia | CTRL vs. SE+1D | 0.0057 | ** | Yes | One-way ANOVA | Tukey's |
| | | | CTRL vs. SE+9D | 0.999 | ns | | | |
| | | | SE+1D vs. SE+9D | 0.006 | $$ | | | |
| | | Mo-mΦ | SE+1D vs. SE+9D | 0.0007 | $$$ | Yes | Unpaired t-test | |
| | | CD11b- | CTRL vs. SE+1D | 0.0417 | * | No | Krukall Wallis | Dunn's |
| | | | CTRL vs. SE+9D | 0.7907 | ns | | | |
| | | | SE+1D vs. SE+9D | 0.4008 | ns | | | |
| | | CTRL | Microglia vs. CD11b- | 0.2000 | ns | No | Mann-Whitney | |
| | | SE+1D | Microglia vs. Mo-mΦ | 0.0188 | # | Yes | RM One-way ANOVA | Tukey's |
| | | | Microglia vs. CD11b- | 0.7902 | ns | | | |
| | | | Mo-mΦ vs. CD11b- | 0.0119 | € | | | |
| | | SE+9D | Microglia vs. Mo-mΦ | 0.4681 | ns | Yes | RM One-way ANOVA | Tukey's |
| | | | Microglia vs. CD11b- | 0.568 | ns | | | |
| | | | Mo-mΦ vs. CD11b- | 0.9771 | ns | | | |
| | MCP1 7W | CTRL 7W vs. SE+7W | Microglia | 0.0011 | ** | Yes | Unpaired t-test | |
| | | | CD11b- | 0.5056 | ns | Yes | Unpaired t-test | |
| | | CTRL 7W | Microglia vs. CD11b- | 0.9695 | ns | Yes | Paired t-test | |
| | | SE+7W | Microglia vs Mo-mΦ | 0.6057 | ns | Yes | RM One-way ANOVA | Tukey's |
| | | | Microglia vs CD11b- | 0.1461 | ns | | | |
| | | | Mo-mΦ vs CD11b- | 0.4178 | ns | | | |
| 4G | IL-10 | Microglia | CTRL vs. SE+1D | 0.6991 | ns | No | Krukall Wallis | Dunn's |
| | | | CTRL vs. SE+9D | 0.0512 | ns | | | |
| | | | SE+1D vs. SE+9D | 0.6991 | ns | | | |
| | | Mo-mΦ | SE+1D vs. SE+9D | 0.0403 | $ | Yes | Unpaired t-test | |
| | | CD11b- | CTRL vs. SE+1D | >0.9999 | ns | No | Krukall Wallis | Dunn's |
| | | | CTRL vs. SE+9D | >0.9999 | ns | | | |
| | | | SE+1D vs. SE+9D | >0.9999 | ns | | | |
| | | CTRL | Microglia vs. CD11b- | 0.4000 | ns | No | Mann-Whitney | |
| | | SE+1D | Microglia vs. Mo-mΦ | 0.0470 | # | Yes | RM One-way ANOVA | Tukey's |
| | | | Microglia vs. CD11b- | 0.9835 | ns | | | |
| | | | Mo-mΦ vs. CD11b- | 0.0381 | € | | | |
| | | SE+9D | Microglia vs. Mo-mΦ | 0.3074 | ns | No | Friedman | Dunn's |
| | | | Microglia vs. CD11b- | >0.9999 | ns | | | |
| | | | Mo-mΦ vs. CD11b- | 0.1237 | ns | | | |

| | | | | | | | | |
|---|---|---|---|---|---|---|---|---|
| | IL-10 7W | CTRL 7W vs. SE+7W | Microglia | 0.4466 | ns | Yes | Unpaired t-test | |
| | | | CD11b- | 0.4000 | ns | No | Mann-Whitney | |
| | | CTRL 7W | Microglia vs. CD11b- | 0.5312 | ns | Yes | Paired t-test | |
| | | SE+7W | Microglia vs Mo-mΦ | >0.9999 | ns | No | Friedman test | Dunn's |
| | | | Microglia vs CD11b- | >0.9999 | ns | | | |
| | | | Mo-mΦ vs CD11b- | 0.8902 | ns | | | |
| 4H | IL-13 | Microglia | CTRL vs. SE+1D | 0.7025 | ns | Yes | One-way ANOVA | Tukey's |
| | | | CTRL vs. SE+9D | 0.3597 | ns | | | |
| | | | SE+1D vs. SE+9D | 0.7914 | ns | | | |
| | | Mo-mΦ | SE+1D vs. SE+9D | 0.0918 | ns | Yes | Unpaired t-test | |
| | | CD11b- | CTRL vs. SE+1D | 0.4702 | ns | No | Krukall Wallis | Dunn's |
| | | | CTRL vs. SE+9D | 0.9977 | ns | | | |
| | | | SE+1D vs. SE+9D | >0.9999 | ns | | | |
| | | CTRL | Microglia vs. CD11b- | 0.8000 | ns | No | Mann-Whitney | |
| | | SE+1D | Microglia vs. Mo-mΦ | 0.1946 | ns | Yes | One-way ANOVA | Tukey's |
| | | | Microglia vs. CD11b- | 0.1517 | ns | | | |
| | | | Mo-mΦ vs. CD11b- | 0.9790 | ns | | | |
| | | SE+9D | Microglia vs. Mo-mΦ | 0.9879 | ns | Yes | One-way ANOVA | Tukey's |
| | | | Microglia vs. CD11b- | 0.4621 | ns | | | |
| | | | Mo-mΦ vs. CD11b- | 0.5392 | ns | | | |
| | IL-13 7W | CTRL 7W vs. SE+7W | Microglia | 0.8574 | ns | Yes | Unpaired t-test | |
| | | | CD11b- | 0.5321 | ns | Yes | Unpaired t-test | |
| | | CTRL 7W | Microglia vs. CD11b- | 0.2847 | ns | Yes | Unpaired t-test | |
| | | SE+7W | Microglia vs Mo-mΦ | 0.9850 | ns | Yes | One-way ANOVA | Tukey's |
| | | | Microglia vs CD11b- | 0.9935 | ns | | | |
| | | | Mo-mΦ vs CD11b- | 0.9594 | ns | | | |
| 4I | Arg1 | Microglia | CTRL vs. SE+1D | 0.4081 | ns | No | Krukall Wallis | Dunn's |
| | | | CTRL vs. SE+9D | 0.8902 | ns | | | |
| | | | SE+1D vs. SE+9D | 0.0338 | $ | | | |
| | | Mo-mΦ | SE+1D vs. SE+9D | 0.0003 | $$$ | Yes | Unpaired t-test | |
| | | CD11b- | CTRL vs. SE+1D | 0.0760 | ns | No | Krukall Wallis | Dunn's |
| | | | CTRL vs. SE+9D | >0.9999 | ns | | | |
| | | | SE+1D vs. SE+9D | 0.2867 | ns | | | |
| | | CTRL | Microglia vs. CD11b- | 0.8000 | ns | No | Mann-Whitney | |
| | | SE+1D | Microglia vs. Mo-mΦ | <0.0001 | #### | Yes | RM One-way ANOVA | Tukey's |
| | | | Microglia vs. CD11b- | 0.9971 | ns | | | |
| | | | Mo-mΦ vs. CD11b- | <0.0001 | €€€€ | | | |
| | | SE+9D | Microglia vs. Mo-mΦ | >0.9999 | ns | No | Friedman | Dunn's |
| | | | Microglia vs. CD11b- | 0.1237 | ns | | | |
| | | | Mo-mΦ vs. CD11b- | 0.3074 | ns | | | |
| | Arg1 7W | CTRL 7W vs. SE+7W | Microglia | 0.0114 | * | Yes | Unpaired t-test | |
| | | | CD11b- | 0.9750 | ns | Yes | Unpaired t-test | |
| | | CTRL 7W | Microglia vs. CD11b- | 0.0243 | # | Yes | Paired t-test | |
| | | SE+7W | Microglia vs Mo-mΦ | >0.9999 | ns | No | Friedman | Dunn's |
| | | | Microglia vs CD11b- | 0.3074 | ns | | | |
| | | | Mo-mΦ vs CD11b- | 0.1237 | ns | | | |
| 4J | CD206 | Microglia | CTRL vs. SE+1D | 0.5391 | ns | No | Krukall Wallis | Dunn's |
| | | | CTRL vs. SE+9D | 0.0219 | * | | | |
| | | | SE+1D vs. SE+9D | 0.5391 | ns | | | |
| | | Mo-mΦ | SE+1D vs. SE+9D | 0.0002 | $$$ | Yes | Unpaired t-test | |
| | | CD11b- | CTRL vs. SE+1D | >0.9999 | ns | No | Krukall Wallis | Dunn's |
| | | | CTRL vs. SE+9D | 0.6991 | ns | | | |
| | | | SE+1D vs. SE+9D | 0.1365 | ns | | | |
| | | CTRL | Microglia vs. CD11b- | 0.2000 | ns | No | Mann-Whitney | |
| | | SE+1D | Microglia vs. Mo-mΦ | <0.0001 | #### | Yes | RM One-way ANOVA | Tukey's |
| | | | Microglia vs. CD11b- | 0.0514 | ns | | | |
| | | | Mo-mΦ vs. CD11b- | <0.0001 | €€€€ | | | |
| | | SE+9D | Microglia vs. Mo-mΦ | 0.3391 | ns | Yes | RM One-way ANOVA | Tukey's |
| | | | Microglia vs. CD11b- | 0.4628 | ns | | | |

| | | | | | | | | |
|---|---|---|---|---|---|---|---|---|
| | | | Mo-mΦ vs. CD11b- | 0.0896 | ns | | | |
| | CD206 7W | CTRL 7W vs. SE+7W | Microglia | 0.5035 | ns | Yes | Unpaired t-test | |
| | | | CD11b- | 0.8760 | ns | Yes | Unpaired t-test | |
| | | CTRL 7W | Microglia vs. CD11b- | 0.2199 | ns | Yes | Paired t-test | |
| | | SE+7W | Microglia vs Mo-mΦ | 0.1185 | ns | Yes | RM One-way ANOVA | Tukey's |
| | | | Microglia vs CD11b- | 0.6198 | ns | | | |
| | | | Mo-mΦ vs CD11b- | 0.0353 | € | | | |

### Figure S2

| | | | p-value | | | | Normality ? | Test | Post-hoc |
|---|---|---|---|---|---|---|---|---|---|
| | | | vs. CTRL (*) | | vs. SE+7h ($) | | | | |
| S2A | MCP1 | SE+7h | <0.0001 | **** | | | Yes | One-way ANOVA | Tukey's |
| | | SE+1D | <0.0001 | **** | <0.0001 | $$$$ | | | |
| | | SE+9D | 0.9997 | ns | <0.0001 | $$$$ | | | |
| | | SE+7W | 0.0009 | *** | | | Yes | Unpaired t-test | |
| S2B | MIP1α | SE+7h | <0.0001 | **** | | | Yes | One-way ANOVA | Tukey's |
| | | SE+1D | 0.3106 | ns | <0.0001 | $$$$ | | | |
| | | SE+9D | 0.8116 | ns | <0.0001 | $$$$ | | | |
| | | SE+7W | <0.0001 | **** | | | Yes | Unpaired t-test | |

### Figure S4

| | | | p-value | | | | Normality ? | Test | Post-hoc |
|---|---|---|---|---|---|---|---|---|---|
| | | | vs. CTRL (*) | | vs. SE+7h ($) | | | | |
| S4A-B | CD68 (HI) | SE+7h | >0.9999 | ns | | | No | Krukall Wallis | Dunn's |
| | | SE+1D | 0.0423 | * | >0.9999 | ns | | | |
| | | SE+9D | <0.0001 | **** | 0.0225 | $ | | | |
| | | SE+7W | <0.0001 | **** | | | Yes | Unpaired t-test | |
| | MCP1 (HI) | SE+7h | <0.0001 | **** | | | Yes | One-way ANOVA | Tukey's |
| | | SE+1D | <0.0001 | **** | <0.0001 | $$$$ | | | |
| | | SE+9D | 0.9997 | ns | <0.0001 | $$$$ | | | |
| | | SE+7W | 0.0009 | *** | | | Yes | Unpaired t-test | |
| | CD68 (VLR) | SE+7h | >0.9999 | ns | | | Yes | One-way ANOVA | Tukey's |
| | | SE+1D | 0.5368 | ns | 0.5988 | ns | | | |
| | | SE+9D | <0.0001 | **** | <0.0001 | $$$$ | | | |
| | | SE+7W | <0.0001 | **** | | | Yes | Unpaired t-test | |
| | MCP1 (VLR) | SE+7h | 0.0210 | * | | | No | Krukall Wallis | Dunn's |
| | | SE+1D | <0.0001 | **** | >0.9999 | ns | | | |
| | | SE+9D | 0.5909 | ns | 0.8838 | ns | | | |
| | | SE+7W | 0.3450 | ns | | | No | Mann-Whitney | |
| | CD68 (ThD) | SE+7h | 0.9991 | ns | | | Yes | One-way ANOVA | Tukey's |
| | | SE+1D | 0.5102 | ns | 0.6294 | ns | | | |
| | | SE+9D | <0.0001 | **** | <0.0001 | $$$$ | | | |
| | | SE+7W | 0.0080 | ** | | | No | Mann-Whitney | |
| | MCP1 (ThD) | SE+7h | 0.0009 | *** | | | Yes | One-way ANOVA | Tukey's |
| | | SE+1D | 0.0031 | ** | 0.8748 | ns | | | |
| | | SE+9D | 0.9999 | ns | 0.0007 | $$$ | | | |
| | | SE+7W | 0.0007 | *** | | | No | Mann-Whitney | |
| | | | | p-value | | | | p-value | |
| | CD68 | CTRL | HI vs. VLR | 0,1381 | ns | SE+9D | HI vs. VLR | 0,0050 | ** |
| | | | HI vs. ThD | 0,6921 | ns | | HI vs. ThD | 0,1197 | ns |
| | | | VLR vs. ThD | 0,2291 | ns | | VLR vs. ThD | 0,0213 | * |
| | | SE+7h | HI vs. VLR | 0,8542 | ns | CTRL+7W | HI vs. VLR | 0,0012 | ** |
| | | | HI vs. ThD | 0,7893 | ns | | HI vs. ThD | 0,0015 | ** |
| | | | VLR vs. ThD | 0,6194 | ns | | VLR vs. ThD | 0,2578 | ns |
| | | SE+1D | HI vs. VLR | 0,3156 | ns | SE+7W | HI vs. VLR | 0,3413 | ns |
| | | | HI vs. ThD | 0,0978 | ns | | HI vs. ThD | 0,0004 | *** |
| | | | VLR vs. ThD | 0,0001 | *** | | VLR vs. ThD | 0,0001 | *** |
| | MCP1 | CTRL | HI vs. VLR | 0,9154 | ns | SE+9D | HI vs. VLR | 0,1732 | ns |

| | | | | | | | | | |
|---|---|---|---|---|---|---|---|---|---|
| | | | HI vs. ThD | 0,3991 | ns | | HI vs. ThD | 0,0444 | * |
| | | | VLR vs. ThD | 0,5263 | ns | | VLR vs. ThD | 0,0014 | ** |
| | | SE+7h | HI vs. VLR | 0,0309 | * | CTRL+7W | HI vs. VLR | 0,1318 | ns |
| | | | HI vs. ThD | 0,0095 | ** | | HI vs. ThD | 0,2956 | ns |
| | | | VLR vs. ThD | 0,7789 | ns | | VLR vs. ThD | 0,0573 | ns |
| | | SE+1D | HI vs. VLR | 0,9854 | ns | SE+7W | HI vs. VLR | 0,7834 | ns |
| | | | HI vs. ThD | 0,0001 | *** | | HI vs. ThD | 0,1332 | ns |
| | | | VLR vs. ThD | 0,0014 | ** | | VLR vs. ThD | 0,3894 | ns |

| | | | p-value | | Normality ? | Test | Post-hoc |
|---|---|---|---|---|---|---|---|
| S4H | CD68+ round cells | Hilus vs. HI | 0.0005 | *** | Yes | One-way ANOVA | Tukey's |
| | | Hilus vs. VLR | 0.0055 | ** | | | |
| | | Hilus vs. ThD | 0.0019 | ** | | | |
| | | HI vs. VLR | 0.6475 | ns | | | |
| | | HI vs. ThD | 0.9116 | ns | | | |
| | | VLR vs. ThD | 0.9523 | ns | | | |

**Figure S5**

| | | | p-value | | | | Normality ? | Test | Post-hoc |
|---|---|---|---|---|---|---|---|---|---|
| | | | vs. CTRL (*) | | vs. SE+7h ($) | | | | |
| S5A | IL-1β (HI) | SE+7h | 0.0001 | *** | | | No | Krukall Wallis | Dunn's |
| | | SE+1D | 0.0065 | ** | >0.9999 | ns | | | |
| | | SE+9D | 0.5909 | ns | 0.0305 | $ | | | |
| | | SE+7W | 0.0029 | ** | | | Yes | Unpaired t-test | |
| S5B | IL-6 (HI) | SE+7h | <0.0001 | **** | | | Yes | One-way ANOVA | Tukey's |
| | | SE+1D | 0.0004 | *** | 0.0009 | $$$ | | | |
| | | SE+9D | >0.9999 | ns | <0.0001 | $$$$ | | | |
| | | SE+7W | 0.7776 | ns | | | No | Mann-Whitney | |
| S5C | TNFα (HI) | SE+7h | <0.0001 | **** | | | Yes | One-way ANOVA | Tukey's |
| | | SE+1D | 0.0014 | ** | 0.0006 | $$$ | | | |
| | | SE+9D | 0.9966 | ns | <0.0001 | $$$$ | | | |
| | | SE+7W | 0.6620 | ns | | | No | Mann-Whitney | |
| S5D | Pro-inflam index (HI) | SE+7h | <0.0001 | **** | | | Yes | One-way ANOVA | Tukey's |
| | | SE+1D | <0.0001 | **** | <0.0001 | $$$$ | | | |
| | | SE+9D | 0.9908 | ns | <0.0001 | $$$$ | | | |
| | | SE+7W | 0.2393 | ns | | | Yes | Unpaired t-test | |
| S5E | IL-4 (HI) | SE+7h | 0.0140 | * | | | No | Krukall Wallis | Dunn's |
| | | SE+1D | 0.0478 | * | >0.9999 | ns | | | |
| | | SE+9D | >0.9999 | ns | 0.0076 | $$ | | | |
| | | SE+7W | 0.6270 | ns | | | Yes | Unpaired t-test | |
| S5F | IL-10 (HI) | SE+7h | <0.0001 | **** | | | Yes | One-way ANOVA | Tukey's |
| | | SE+1D | 0.0060 | ** | 0.1649 | ns | | | |
| | | SE+9D | 0.2276 | ns | 0.0030 | $$ | | | |
| | | SE+7W | 0.6217 | ns | | | Yes | Unpaired t-test | |
| S5G | IL-13 (HI) | SE+7h | 0.0819 | ns | | | Yes | One-way ANOVA | Tukey's |
| | | SE+1D | 0.0813 | ns | 0.9994 | ns | | | |
| | | SE+9D | 0.9973 | ns | 0.0475 | $ | | | |
| | | SE+7W | 0.0565 | ns | | | Yes | Unpaired t-test | |
| S5H | Anti-inflam index (HI) | SE+7h | 0.0003 | *** | | | Yes | One-way ANOVA | Tukey's |
| | | SE+1D | 0.0078 | ** | 0.3852 | ns | | | |
| | | SE+9D | 0.9606 | ns | 0.0005 | $$$ | | | |
| | | SE+7W | 0.3047 | ns | | | Yes | Unpaired t-test | |
| S5A | IL-1β (VLR) | SE+7h | 0.0001 | *** | | | Yes | One-way ANOVA | Tukey's |
| | | SE+1D | <0.0001 | **** | 0.9982 | ns | | | |
| | | SE+9D | 0.9112 | ns | 0.0003 | $$$ | | | |
| | | SE+7W | 0.0357 | * | | | Yes | Unpaired t-test | |
| S5B | IL-6 (VLR) | SE+7h | <0.0001 | **** | | | Yes | One-way ANOVA | Tukey's |
| | | SE+1D | <0.0001 | **** | 0.9976 | ns | | | |
| | | SE+9D | >0.9999 | ns | <0.0001 | $$$$ | | | |

| | | | | | | | | | |
|---|---|---|---|---|---|---|---|---|---|
| | | SE+7W | <0.0001 | **** | | | Yes | Unpaired t-test | |
| S5C | TNFα (VLR) | SE+7h | 0.0038 | ** | | | No | Krukall Wallis | Dunn's |
| | | SE+1D | 0.1193 | ns | >0.9999 | ns | | | |
| | | SE+9D | >0.9999 | ns | 0.0018 | $$ | | | |
| | | SE+7W | 0.6620 | ns | | | No | Mann-Whitney | |
| S5D | Pro-inflam index (VLR) | SE+7h | 0.0046 | ** | | | No | Krukall Wallis | Dunn's |
| | | SE+1D | 0.0049 | ** | >0.9999 | ns | | | |
| | | SE+9D | >0.9999 | ns | 0.0591 | ns | | | |
| | | SE+7W | 0.1419 | ns | | | No | Mann-Whitney | |
| S5E | IL-4 (VLR) | SE+7h | <0.0001 | **** | | | Yes | One-way ANOVA | Tukey's |
| | | SE+1D | 0.7237 | ns | <0.0001 | $$$$ | | | |
| | | SE+9D | >0.9999 | ns | <0.0001 | $$$$ | | | |
| | | SE+7W | 0.9964 | ns | | | Yes | Unpaired t-test | |
| S5F | IL-10 (VLR) | SE+7h | 0.0214 | * | | | No | Krukall Wallis | Dunn's |
| | | SE+1D | 0.0019 | ** | >0.9999 | ns | | | |
| | | SE+9D | >0.9999 | ns | 0.1409 | ns | | | |
| | | SE+7W | 0.0027 | ** | | | No | Mann-Whitney | |
| S5G | IL-13 (VLR) | SE+7h | >0.9999 | ns | | | No | Krukall Wallis | Dunn's |
| | | SE+1D | >0.9999 | ns | 0.1326 | ns | | | |
| | | SE+9D | 0.0884 | ns | 0.0014 | $$ | | | |
| | | SE+7W | 0.0003 | *** | | | Yes | Unpaired t-test | |
| S5H | Anti-inflam index (VLR) | SE+7h | 0.0304 | * | | | No | Krukall Wallis | Dunn's |
| | | SE+1D | 0.6681 | ns | >0.9999 | ns | | | |
| | | SE+9D | 0.9402 | ns | 0.0001 | $$$ | | | |
| | | SE+7W | 0.1812 | ns | | | No | Mann-Whitney | |
| S5A | IL-1β (ThD) | SE+7h | 0.0009 | *** | | | No | Krukall Wallis | Dunn's |
| | | SE+1D | 0.0049 | ** | >0.9999 | ns | | | |
| | | SE+9D | >0.9999 | ns | 0.0483 | $ | | | |
| | | SE+7W | 0.0499 | * | | | Yes | Unpaired t-test | |
| S5B | IL-6 (ThD) | SE+7h | <0.0001 | **** | | | Yes | One-way ANOVA | Tukey's |
| | | SE+1D | 0.3651 | ns | <0.0001 | $$$$ | | | |
| | | SE+9D | >0.9999 | ns | <0.0001 | $$$$ | | | |
| | | SE+7W | <0.0001 | **** | | | Yes | Unpaired t-test | |
| S5C | TNFα (ThD) | SE+7h | 0.0431 | * | | | No | Krukall Wallis | Dunn's |
| | | SE+1D | 0.8478 | ns | >0.9999 | ns | | | |
| | | SE+9D | 0.5891 | ns | <0.0001 | $$$$ | | | |
| | | SE+7W | 0.5728 | ns | | | No | Mann-Whitney | |
| S5D | Pro-inflam index (ThD) | SE+7h | <0.0001 | **** | | | Yes | One-way ANOVA | Tukey's |
| | | SE+1D | 0.0048 | ** | <0.0001 | $$$$ | | | |
| | | SE+9D | 0.9998 | ns | <0.0001 | $$$$ | | | |
| | | SE+7W | 0.2284 | ns | | | No | Mann-Whitney | |
| S5E | IL-4 (ThD) | SE+7h | 0.0363 | * | | | No | Krukall Wallis | Dunn's |
| | | SE+1D | >0.9999 | ns | 0.0018 | $$ | | | |
| | | SE+9D | >0.9999 | ns | 0.1561 | ns | | | |
| | | SE+7W | 0.3980 | ns | | | Yes | Unpaired t-test | |
| S5F | IL-10 (ThD) | SE+7h | 0.0004 | *** | | | Yes | One-way ANOVA | Tukey's |
| | | SE+1D | 0.1691 | ns | 0.0361 | $ | | | |
| | | SE+9D | 0.9896 | ns | 0.0005 | $$$ | | | |
| | | SE+7W | 0.2177 | ns | | | Yes | Unpaired t-test | |
| S5G | IL-13 (ThD) | SE+7h | <0.0001 | **** | | | Yes | One-way ANOVA | Tukey's |
| | | SE+1D | 0.4405 | ns | <0.0001 | $$$$ | | | |
| | | SE+9D | >0.9999 | ns | <0.0001 | $$$$ | | | |
| | | SE+7W | 0.1462 | ns | | | Yes | Unpaired t-test | |
| S5H | Anti-inflam index (ThD) | SE+7h | <0.0001 | **** | | | Yes | One-way ANOVA | Tukey's |
| | | SE+1D | 0.0986 | ns | 0.0003 | $$$ | | | |
| | | SE+9D | 0.9886 | ns | <0.0001 | $$$$ | | | |
| | | SE+7W | 0.4333 | ns | | | Yes | Unpaired t-test | |

| | | | | p-value | | | | p-value | |
|---|---|---|---|---|---|---|---|---|---|
| S5A | IL-1β | CTRL | HI vs. VLR | 0.8543 | ns | SE+9D | HI vs. VLR | 0.8081 | ns |
| | | | HI vs. ThD | 0.4351 | ns | | HI vs. ThD | 0.0058 | ** |
| | | | VLR vs. ThD | 0.3433 | ns | | VLR vs. ThD | 0.1475 | ns |
| | | SE+7h | HI vs. VLR | 0.3240 | ns | CTRL+7W | HI vs. VLR | 0.2911 | ns |
| | | | HI vs. ThD | 0.8829 | ns | | HI vs. ThD | 0.1244 | ns |
| | | | VLR vs. ThD | 0.2537 | ns | | VLR vs. ThD | 0.3823 | ns |
| | | SE+1D | HI vs. VLR | 0.0028 | ** | SE+7W | HI vs. VLR | 0.0927 | ns |
| | | | HI vs. ThD | 0.8328 | ns | | HI vs. ThD | 0.0229 | * |
| | | | VLR vs. ThD | 0.0016 | ** | | VLR vs. ThD | 0.3263 | ns |
| S5B | IL-6 | CTRL | HI vs. VLR | 0.0731 | ns | SE+9D | HI vs. VLR | 0.0319 | * |
| | | | HI vs. ThD | 0.0731 | ns | | HI vs. ThD | 0.0319 | * |
| | | | VLR vs. ThD | >0.9999 | ns | | VLR vs. ThD | >0.9999 | ns |
| | | SE+7h | HI vs. VLR | 0.1625 | ns | CTRL+7W | HI vs. VLR | <0.0001 | **** |
| | | | HI vs. ThD | 0.2908 | ns | | HI vs. ThD | <0.0001 | **** |
| | | | VLR vs. ThD | 0.5647 | ns | | VLR vs. ThD | 0.0020 | ** |
| | | SE+1D | HI vs. VLR | 0.0035 | ** | SE+7W | HI vs. VLR | <0.0001 | **** |
| | | | HI vs. ThD | 0.0622 | ns | | HI vs. ThD | <0.0001 | **** |
| | | | VLR vs. ThD | 0.0006 | *** | | VLR vs. ThD | <0.0001 | **** |
| S5C | TNFα | CTRL | HI vs. VLR | 0.6563 | ns | SE+9D | HI vs. VLR | 0.9966 | ns |
| | | | HI vs. ThD | 0.5427 | ns | | HI vs. ThD | 0.0206 | * |
| | | | VLR vs. ThD | 0.9795 | ns | | VLR vs. ThD | 0.0652 | ns |
| | | SE+7h | HI vs. VLR | 0.4244 | ns | CTRL+7W | HI vs. VLR | 0.2090 | ns |
| | | | HI vs. ThD | 0.6157 | ns | | HI vs. ThD | 0.0314 | * |
| | | | VLR vs. ThD | 0.1024 | ns | | VLR vs. ThD | 0.0153 | * |
| | | SE+1D | HI vs. VLR | 0.9945 | ns | SE+7W | HI vs. VLR | 0.0522 | ns |
| | | | HI vs. ThD | 0.8233 | ns | | HI vs. ThD | 0.0314 | * |
| | | | VLR vs. ThD | 0.5462 | ns | | VLR vs. ThD | 0.5089 | ns |
| S5D | Pro-inflam index | CTRL | HI vs. VLR | 0.3044 | ns | SE+9D | HI vs. VLR | 0.5282 | ns |
| | | | HI vs. ThD | 0.8517 | ns | | HI vs. ThD | 0.0068 | ** |
| | | | VLR vs. ThD | 0.3413 | ns | | VLR vs. ThD | 0.1475 | ns |
| | | SE+7h | HI vs. VLR | 0.2236 | ns | CTRL+7W | HI vs. VLR | 0.0241 | * |
| | | | HI vs. ThD | 0.8097 | ns | | HI vs. ThD | 0.0227 | * |
| | | | VLR vs. ThD | 0.2081 | ns | | VLR vs. ThD | 0.4148 | ns |
| | | SE+1D | HI vs. VLR | 0.0028 | ** | SE+7W | HI vs. VLR | 0.0288 | * |
| | | | HI vs. ThD | 0.0553 | ns | | HI vs. ThD | 0.0389 | * |
| | | | VLR vs. ThD | 0.0006 | *** | | VLR vs. ThD | 0.3324 | ns |
| S5E | IL-4 | CTRL | HI vs. VLR | 0.8531 | ns | SE+9D | HI vs. VLR | 0.5988 | ns |
| | | | HI vs. ThD | 0.0388 | * | | HI vs. ThD | 0.0847 | ns |
| | | | VLR vs. ThD | 0.0169 | * | | VLR vs. ThD | 0.0047 | ** |
| | | SE+7h | HI vs. VLR | 0.0070 | ** | CTRL+7W | HI vs. VLR | 0.9671 | ns |
| | | | HI vs. ThD | 0.1171 | ns | | HI vs. ThD | 0.0015 | ** |
| | | | VLR vs. ThD | 0.0100 | ** | | VLR vs. ThD | 0.0042 | ** |
| | | SE+1D | HI vs. VLR | 0.5227 | ns | SE+7W | HI vs. VLR | 0.5898 | ns |
| | | | HI vs. ThD | 0.1134 | ns | | HI vs. ThD | 0.0154 | * |
| | | | VLR vs. ThD | 0.0004 | *** | | VLR vs. ThD | 0.0040 | ** |
| S5F | IL-10 | CTRL | HI vs. VLR | 0.1956 | ns | SE+9D | HI vs. VLR | 0.4647 | ns |
| | | | HI vs. ThD | 0.6124 | ns | | HI vs. ThD | 0.2459 | ns |
| | | | VLR vs. ThD | 0.2305 | ns | | VLR vs. ThD | 0.1225 | ns |
| | | SE+7h | HI vs. VLR | 0.3529 | ns | CTRL+7W | HI vs. VLR | 0.6801 | ns |
| | | | HI vs. ThD | 0.9561 | ns | | HI vs. ThD | 0.9557 | ns |
| | | | VLR vs. ThD | 0.2543 | ns | | VLR vs. ThD | 0.6618 | ns |
| | | SE+1D | HI vs. VLR | 0.0234 | * | SE+7W | HI vs. VLR | 0.0854 | ns |
| | | | HI vs. ThD | 0.5935 | ns | | HI vs. ThD | 0.0903 | ns |
| | | | VLR vs. ThD | 0.0111 | * | | VLR vs. ThD | 0.0471 | * |
| S5G | IL-13 | CTRL | HI vs. VLR | 0.0180 | * | SE+9D | HI vs. VLR | 0.5598 | ns |
| | | | HI vs. ThD | 0.0003 | *** | | HI vs. ThD | 0.0596 | ns |
| | | | VLR vs. ThD | 0.0068 | ** | | VLR vs. ThD | 0.0208 | * |
| | | SE+7h | HI vs. VLR | 0.0897 | ns | CTRL+7W | HI vs. VLR | 0.0053 | ** |

| | | | | | | | | | |
|---|---|---|---|---|---|---|---|---|---|
| | | | HI vs. ThD | 0.1353 | ns | | HI vs. ThD | 0.0087 | ** |
| | | | VLR vs. ThD | 0.0170 | * | | VLR vs. ThD | 0.0028 | ** |
| | | SE+1D | HI vs. VLR | 0.8716 | ns | SE+7W | HI vs. VLR | 0.7980 | ns |
| | | | HI vs. ThD | 0.0380 | * | | HI vs. ThD | 0.0078 | ** |
| | | | VLR vs. ThD | 0.0077 | ** | | VLR vs. ThD | 0.0013 | ** |
| S5H | Anti-inflam Index | CTRL | HI vs. VLR | 0.0082 | ** | SE+9D | HI vs. VLR | 0.7953 | ns |
| | | | HI vs. ThD | <0.0001 | **** | | HI vs. ThD | 0.0616 | ns |
| | | | VLR vs. ThD | 0.0014 | ** | | VLR vs. ThD | 0.0133 | * |
| | | SE+7h | HI vs. VLR | 0.0010 | *** | CTRL+7W | HI vs. VLR | 0.0434 | * |
| | | | HI vs. ThD | 0.0608 | ns | | HI vs. ThD | 0.0098 | ** |
| | | | VLR vs. ThD | 0.0004 | *** | | VLR vs. ThD | 0.0048 | ** |
| | | SE+1D | HI vs. VLR | 0.7901 | ns | SE+7W | HI vs. VLR | 0.1789 | ns |
| | | | HI vs. ThD | 0.0409 | * | | HI vs. ThD | 0.0062 | ** |
| | | | VLR vs. ThD | 0.0022 | ** | | VLR vs. ThD | 0.0005 | *** |

**Figure S6**

| | | | p-value | | | | Normality ? | Test | Post-hoc |
|---|---|---|---|---|---|---|---|---|---|
| | | | vs. CTRL (*) | | vs. SE+1D ($) | | | | |
| S6F | Monocytes/Mo-mΦ | SE+1D | 0.0001 | *** | | | Yes | One-way ANOVA | Tukey's |
| | | SE+9D | 0.0159 | * | 0.0006 | $$$ | | | |
| | | SE+7W | 0.0107 | * | | | Yes | Unpaired t-test | |

**Figure S7**

| | | | | p-value | | Normality ? | Test | Post-hoc |
|---|---|---|---|---|---|---|---|---|
| S7A | Iba1 | Microglia | CTRL vs. SE+1D | 0.0550 | ns | Yes | One-way ANOVA | Tukey's |
| | | | CTRL vs. SE+9D | 0.2051 | ns | | | |
| | | | SE+1D vs. SE+9D | 0.5818 | ns | | | |
| | | Mo-mΦ | SE+1D vs. SE+9D | 0.0773 | ns | Yes | Unpaired t-test | |
| | | CD11b- | CTRL vs. SE+1D | 0.7907 | ns | No | Krukall Wallis | Dunn's |
| | | | CTRL vs. SE+9D | 0.0417 | * | | | |
| | | | SE+1D vs. SE+9D | 0.4008 | ns | | | |
| | | CTRL | Microglia vs. CD11b- | 0.2000 | ns | No | Mann-Whitney | |
| | | SE+1D | Microglia vs. Mo-mΦ | 0.0073 | ## | Yes | One-way ANOVA | Tukey's |
| | | | Microglia vs. CD11b- | 0.0002 | ### | | | |
| | | | Mo-mΦ vs. CD11b- | 0.0069 | €€ | | | |
| | | SE+9D | Microglia vs. Mo-mΦ | 0.8953 | ns | Yes | One-way ANOVA | Tukey's |
| | | | Microglia vs. CD11b- | 0.0635 | ns | | | |
| | | | Mo-mΦ vs. CD11b- | 0.0364 | € | | | |
| | Iba1 7W | CTRL 7W vs. SE+7W | Microglia | 0.7381 | ns | Yes | Unpaired t-test | |
| | | | CD11b- | 0.2000 | ns | No | Mann-Whitney | |
| | | CTRL 7W | Microglia vs. CD11b- | 0.1000 | ns | No | Mann-Whitney | |
| | | SE+7W | Microglia vs Mo-mΦ | 0.0112 | # | Yes | One-way ANOVA | Tukey's |
| | | | Microglia vs CD11b- | 0.0012 | ## | | | |
| | | | Mo-mΦ vs CD11b- | 0.1130 | ns | | | |
| S7B | CD68 | Microglia | CTRL vs. SE+1D | 0.1294 | ns | Yes | One-way ANOVA | Dunn's |
| | | | CTRL vs. SE+9D | 0.0137 | * | | | |
| | | | SE+1D vs. SE+9D | 0.2268 | ns | | | |
| | | Mo-mΦ | SE+1D vs. SE+9D | 0.8410 | ns | Yes | Unpaired t-test | |
| | | CD11b- | CTRL vs. SE+1D | 0.7907 | ns | No | Krukall Wallis | Dunn's |
| | | | CTRL vs. SE+9D | 0.0417 | * | | | |
| | | | SE+1D vs. SE+9D | 0.4008 | ns | | | |
| | | CTRL | Microglia vs. CD11b- | 0.2000 | ns | No | Mann-Whitney | |
| | | SE+1D | Microglia vs. Mo-mΦ | 0.0093 | ## | Yes | RM One-way ANOVA | Tukey's |
| | | | Microglia vs. CD11b- | 0.0676 | ns | | | |
| | | | Mo-mΦ vs. CD11b- | 0.0018 | €€ | | | |
| | | SE+9D | Microglia vs. Mo-mΦ | >0.9999 | ns | No | Friedman | Dunn's |
| | | | Microglia vs. CD11b- | 0.3074 | ns | | | |

| | | | | | | | | |
|---|---|---|---|---|---|---|---|---|
| | | | Mo-mΦ vs. CD11b- | 0.1237 | ns | | | |
| | CD68 7W | CTRL 7W vs. SE+7W | Microglia | 0.0307 | * | Yes | Unpaired t-test | |
| | | | CD11b- | >0.9999 | ns | No | Mann-Whitney | |
| | | CTRL 7W | Microglia vs. CD11b- | 0.2500 | # | No | Wilcoxon Paired | |
| | | SE+7W | Microglia vs Mo-mΦ | >0.9999 | ns | No | Friedman | Dunn's |
| | | | Microglia vs CD11b- | 0.3074 | ns | | | |
| | | | Mo-mΦ vs CD11b- | 0.1237 | ns | | | |
| S7C | CD11b | Microglia | CTRL vs. SE+1D | 0.3480 | ns | No | Krukall Wallis | Dunn's |
| | | | CTRL vs. SE+9D | >0.9999 | ns | | | |
| | | | SE+1D vs. SE+9D | 0.0405 | $ | | | |
| | | Mo-mΦ | SE+1D vs. SE+9D | 0.1649 | ns | Yes | Unpaired t-test | |
| | | CD11b- | CTRL vs. SE+1D | 0.7907 | ns | No | Krukall Wallis | Dunn's |
| | | | CTRL vs. SE+9D | 0.0417 | * | | | |
| | | | SE+1D vs. SE+9D | 0.4008 | ns | | | |
| | | CTRL | Microglia vs. CD11b- | 0.2000 | ns | No | Mann-Whitney | |
| | | SE+1D | Microglia vs. Mo-mΦ | <0.0001 | #### | Yes | RM One-way ANOVA | Tukey's |
| | | | Microglia vs. CD11b- | <0.0001 | #### | | | |
| | | | Mo-mΦ vs. CD11b- | 0.0077 | €€ | | | |
| | | SE+9D | Microglia vs. Mo-mΦ | >0.9999 | ns | No | Friedman | Dunn's |
| | | | Microglia vs. CD11b- | 0.3074 | ns | | | |
| | | | Mo-mΦ vs. CD11b- | 0.1237 | ns | | | |
| | CD11b 7W | CTRL 7W vs. SE+7W | Microglia | 0.8717 | ns | Yes | Unpaired t-test | |
| | | | CD11b- | 0.4829 | ns | Yes | Unpaired t-test | |
| | | CTRL 7W | Microglia vs. CD11b- | 0.0408 | # | Yes | Paired t-test | |
| | | SE+7W | Microglia vs Mo-mΦ | 0.0701 | ns | Yes | RM One-way ANOVA | Tukey's |
| | | | Microglia vs CD11b- | 0.014 | # | | | |
| | | | Mo-mΦ vs CD11b- | 0.219 | ns | | | |
| S7D | CD11a | Microglia | CTRL vs. SE+1D | 0.0008 | *** | Yes | One-way ANOVA | Dunn's |
| | | | CTRL vs. SE+9D | 0.1024 | ns | | | |
| | | | SE+1D vs. SE+9D | 0.0067 | $$ | | | |
| | | Mo-mΦ | SE+1D vs. SE+9D | 0.8789 | ns | Yes | Unpaired t-test | |
| | | CD11b- | CTRL vs. SE+1D | 0.7907 | ns | No | Krukall Wallis | Dunn's |
| | | | CTRL vs. SE+9D | 0.0417 | * | | | |
| | | | SE+1D vs. SE+9D | 0.4008 | ns | | | |
| | | CTRL | Microglia vs. CD11b- | 0.2000 | ns | No | Mann-Whitney | |
| | | SE+1D | Microglia vs. Mo-mΦ | 0.1647 | ns | Yes | RM One-way ANOVA | Tukey's |
| | | | Microglia vs. CD11b- | 0.0006 | ### | | | |
| | | | Mo-mΦ vs. CD11b- | 0.0014 | €€ | | | |
| | | SE+9D | Microglia vs. Mo-mΦ | 0.3606 | ns | Yes | RM One-way ANOVA | Tukey's |
| | | | Microglia vs. CD11b- | 0.2104 | ns | | | |
| | | | Mo-mΦ vs. CD11b- | 0.0471 | € | | | |
| | CD11a 7W | CTRL 7W vs. SE+7W | Microglia | 0.3848 | ns | Yes | Unpaired t-test | |
| | | | CD11b- | 0.8428 | ns | Yes | Unpaired t-test | |
| | | CTRL 7W | Microglia vs. CD11b- | 0.0467 | # | Yes | Paired t-test | |
| | | SE+7W | Microglia vs Mo-mΦ | 0.7471 | ns | Yes | RM One-way ANOVA | Tukey's |
| | | | Microglia vs CD11b- | 0.105 | ns | | | |
| | | | Mo-mΦ vs CD11b- | 0.2262 | ns | | | |
